# Sir3 Heterochromatin Protein Promotes NHEJ by Direct Inhibition of Sae2

**DOI:** 10.1101/2021.05.26.445723

**Authors:** Hélène Bordelet, Rafaël Costa, Clémentine Brocas, Jordane Dépagne, Xavier Veaute, Didier Busso, Amandine Batté, Raphaël Guérois, Stéphane Marcand, Karine Dubrana

## Abstract

Heterochromatin is a conserved feature of eukaryotic chromosomes, with central roles in gene expression regulation and maintenance of genome stability. How heterochromatin proteins regulate DNA repair remains poorly described. In *Saccharomyces cerevisiae*, the Silent Information Regulator (SIR) complex assembles heterochromatin-like chromatin at subtelomeres. SIR-mediated repressive chromatin limits double strand break (DSB) resection protecting damaged chromosome ends during HR. As resection initiation marks the cross-road between repair by non-homologous end joining (NHEJ) or HR, we asked whether SIR-mediated heterochromatin regulates NHEJ. We show that SIRs promote NHEJ through two pathways, one depending on repressive chromatin assembly, and the other relying on Sir3 in a manner that is independent of its heterochromatin-promoting function. Sir3 physically interacts with Sae2 and this interaction impairs Sae2-dependent MRX functions. As a consequence, Sir3 limits Mre11-mediated resection, delays MRX removal from DSB ends and promotes NHEJ.

## Introduction

DNA double strand breaks (DSBs) are genotoxic lesions typically repaired by two conserved repair pathways: Non-Homologous End Joining (NHEJ) and Homologous Recombination (HR). NHEJ ligates DSB ends with minimal or no processing, and acts throughout the cell cycle. Repair by HR requires a homologous template for repair, the resection of the DSB ends, and occurs in S and G2 phases. Initiation of DSB resection thus represents a decision point between NHEJ and HR, upon which various cellular inputs converge.

DSB ends are rapidly bound by the Ku70/80 and Mre11-Rad50-Xrs2^NBS1^ (MRX^MRN^) end binding complexes. In *S. cerevisiae*, both complexes aid recruitment of the NHEJ ligation complex composed of the yeast DNA ligase IV Dnl4 (Lig4) and its XRCC4/XLF-like regulatory subunits Lif1 and Nej1 (Palmbos et al. 2005, 2008; Matsuzaki et al. 2008; Chen and Tomkinson 2011; Mahaney et al. 2014).

In addition to its function in NHEJ, the MRX^MRN^ complex is key to shifting repair towards HR when stimulated to initiate resection by Sae2. Indeed, Sae2 activates the endonuclease activity of MRX^MRN^, which cleaves the 5’ strand of the DSB end (Cannavo and Cejka 2014; Bazzano et al. 2021). This provides an entry point for MRX 3’-5’ exonuclease activity, which degrades the DNA towards the DSB, creating a short ssDNA extensions that can no longer be ligated by the canonical NHEJ machinery (Mimitou and Symington 2008; Garcia et al. 2011; Cannavo and Cejka 2014). Impairment of Sae2-MRX dependent resection increases error-prone NHEJ, highlighting the role of Sae2 in coordinating DSB repair pathway choice (Lee and Lee 2007; Huertas et al. 2008).

As a key determinant of NHEJ/HR repair balance, Sae2 activity and protein levels are tightly regulated. Sae2 activity is cell cycle regulated and restricted to S-G2 by CDK- dependent phosphorylation (Huertas et al. 2008). Upon DNA damage, the Tel1 and Mec1 checkpoint kinases phosphorylate Sae2, altering its oligomerization state and forming units active for repair (Baroni et al. 2004; Fu et al. 2014). Sae2 is also negatively regulated by acetylation, which favours its degradation by autophagy thus preventing the persistence of active Sae2 in the cell (Robert et al. 2011; Fu et al. 2014).

In cells, DSB repair does not occur on naked DNA, but in the context of chromatin, which modulates repair efficiency and outcome in several organisms (Goodarzi et al. 2008; Chiolo et al. 2011; Lemaître et al. 2014; Tsouroula et al. 2016; Batté et al. 2017). In *S. cerevisiae* haploid cells, heterochromatin-like chromatin (also called silent chromatin) establishes at the two cryptic mating type loci (*HM* loci) and at each of the 32 subtelomeric loci. Its core components are histone H4 lysine 16 deacetylated nucleosomes, which are bridged by the histone-binding factor Sir3 in complex with the protein Sir4 and the histone deacetylase Sir2 (Behrouzi et al. 2016; Gartenberg and Smith 2016; Faure et al. 2019). Sir2 deacetylates histone H4 lysine 16 thus promoting Sir3 binding and propagation along chromatin. The limiting factor of heterochromatin propagation is Sir3, and its overexpression is sufficient to increase silent chromatin spreading and transcriptional repression in subtelomeric regions, providing an ideal genetic tool to modulate silent chromatin at given sites (Renauld et al. 1993; Hecht et al. 1996; Strahl-Bolsinger et al. 1997; Katan-Khaykovich and Struhl 2005). Sir3 can be seen as the functional ortholog of the heterochromatin factor HP1 that binds histones H3 methylated on lysine 9 in other eukaryotes (Larson et al. 2017; Strom et al. 2017; Machida et al. 2018; Allshire and Madhani 2018). In addition, general heterochromatin properties are conserved in budding yeast such as *cis* and *trans* cooperativity in the establishment of transcription repressive compartments, clustering at the nuclear periphery and near the nucleolus, epigenetic variegation and late replication initiation (Meister and Taddei 2013; Ruault et al. 2021).

SIR proteins also contribute to genome stability in several ways. Sir4 inhibits telomere end fusions by NHEJ (Marcand et al. 2008) and favours telomere elongation through telomerase recruitment (Dalby et al. 2013; Hass and Zappulla 2015; Chen et al. 2018). However, the SIR complex also indirectly promotes NHEJ, as derepression of the *HM* loci in *sir* mutants and the resulting expression of the a1-alpha2 repressor inhibits NHEJ through negative transcriptional regulation of Nej1, and to a lesser extent Lif1 (Aström et al. 1999; Lee et al. 1999; Kegel et al. 2001; Frank-Vaillant and Marcand 2001; Valencia et al. 2001). Finally, we recently showed that SIR-mediated heterochromatin structure protects subtelomeric DSBs from extensive resection (Batté et al. 2017). Whether SIR proteins also inhibit resection initiation and as such play a direct NHEJ-promoting role at subtelomeres is unknown.

Here we found that Sir3 promotes NHEJ in *cis* through heterochromatin formation, as well as in *trans* independently of heterochromatin formation. The *trans* effect relies on a direct interaction between Sir3 and Sae2 that regulates NHEJ repair. This interaction, between the Sir3 conserved AAA+ domain and the C-terminal domain of Sae2, inhibits Sae2 functions. Sae2-Sir3 interaction limits Sae2-MRX dependent resection and favours NHEJ. This function is separable from Sir3-mediated heterochromatin assembly, revealing a new role for SIRs in regulating DSB repair. Sir3 not only promotes genome stability as part of heterochromatin, but is also a direct negative regulator of Sae2, and thus a pro-NHEJ repair factor.

## Results

### NHEJ is increased in *cis* and in *trans* by Sir3 overexpression

Yeast heterochromatin (Sir3-mediated silent chromatin) delays DSB resection, favouring accurate repair by HR near chromosome ends (Batté et al. 2017). Nevertheless, heterochromatin impact on NHEJ has not been addressed. To explore this issue, we used erroneous NHEJ repair of an I-SceI-induced DSB as a proxy for NHEJ efficiency. To establish heterochromatin at the I-SceI site, we exploited the ability of Sir3 overexpression to spread heterochromatin specifically along subtelomeric regions (Batté et al. 2017). The I-SceI site inserted at a subtelomere (1.4 kb from *TEL6R*) is embedded in euchromatin in wild-type (WT) cells but assembled in heterochromatin in cells overexpressing Sir3. Conversely, the I-SceI site inserted at an intrachromosomic position (*LYS2* locus, 300 kb from the closest telomere) remains euchromatic in both contexts (Hocher et al. 2018). Continuous I-SceI expression, driven by a galactose-inducible promoter, is lethal unless NHEJ repairs the DSB with a sequence change that prevents a new cleavage by I-SceI (Fig 1A). Survival frequency was around 10^-3^ in WT cells, and was reduced 10-fold in cells lacking Ligase 4 (Dnl4), indicating that most events leading to survival were products of classical NHEJ (Fig 1B, 1C).

**Figure 1:**
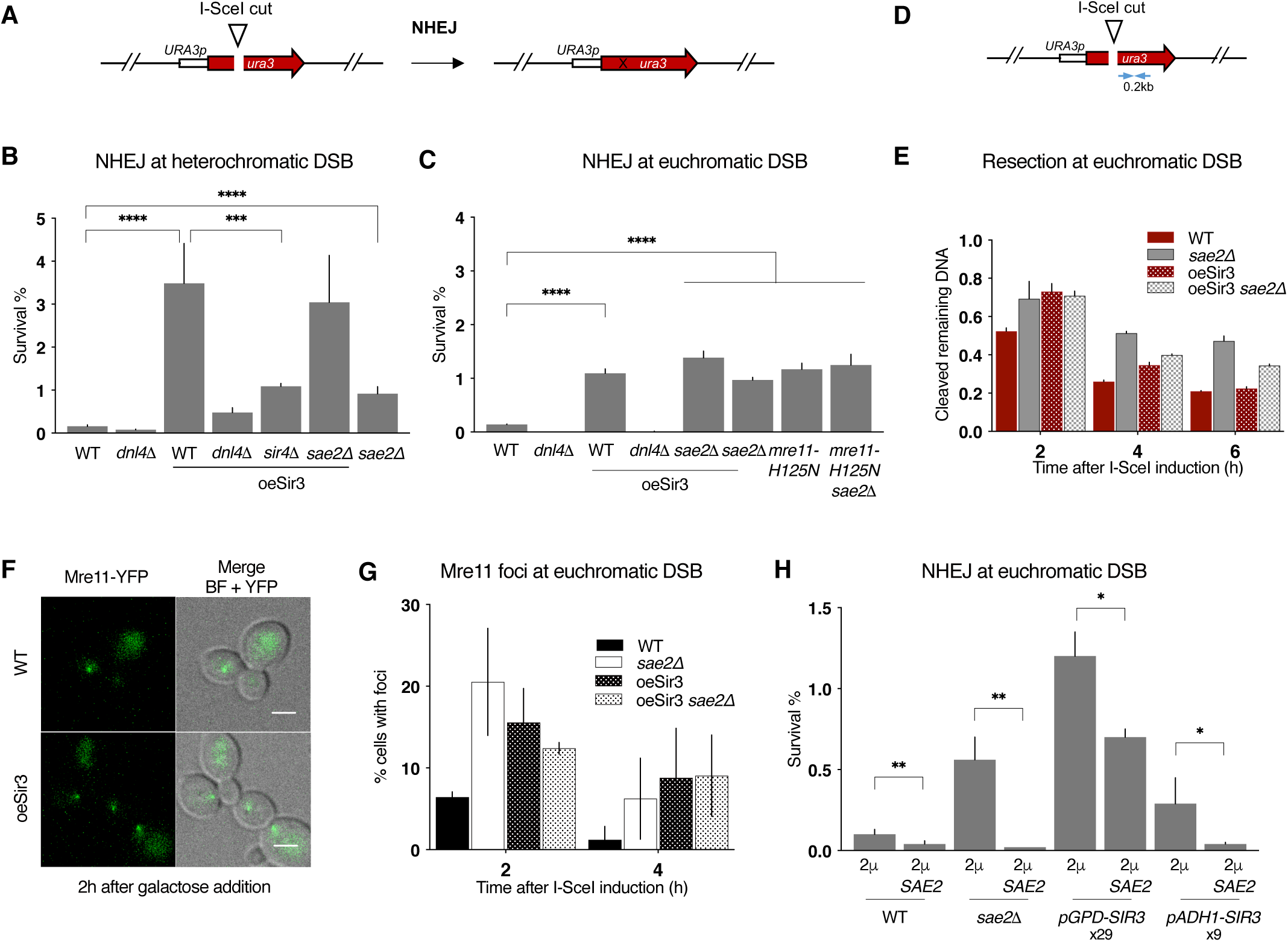
**Sir3 overexpression inhibits Sae2 and increases error-prone NHEJ** A. Schematic representation of the assay used to estimate error-prone NHEJ at euchromatic DSB. B. Survival frequencies observed after DSB induction at TEL6R in WT, dnl4Δ sir4Δ cells, expressing or not high levels of Sir3p (oeSir3 and WT respectively). Error bars indicate survival standard error (SEM) of at least three independent experiments. C. Survival frequencies observed after DSB induction at LYS2 in the indicated strains. Error bars indicate survival standard error (SEM) of at least three independent experiments. D. Schematic representation of the LYS2 locus with primers located at 1 kb from the I-SceI site for the DNA measurements (blue arrows). E. DNA levels measured at 0.2 kb from the I-SceI cut site at LYS2 after 2h DSB induction by qPCR in WT and sae2Δ cells expressing or not high levels of Sir3p (oeSir3 and WT respectively). DNA levels were normalized to DNA levels at the OGG1 locus and corrected for differences in DSB cleavage efficiency (see Materials and Methods for details). Error bars represent the standard deviation (SD) of three independent experiments. F. Representative images of Mre11-YFP foci in response to an I-SceI-induced DSB at LYS2 in WT cells, expressing or not high levels of Sir3p (oeSir3 and WT respectively). Scale bars are 2 μm G. Quantification of cells with DSB induced Mre11-YFP foci after DSB induction at LYS2 I-SceI cleavage site in WT, sae2Δ and Sir3 overexpressing (oeSir3) strains. Error bars indicate survival standard error (SEM) of at least three independent experiments. H. Survival frequencies after DSB induction at LYS2 locus, in strains where SIR3 is expressed from its native, pADH1 or pGPD promoters respectively and in which SAE2 is expressed or not from a high copy number 2µ plasmid. Fold increase in Sir3 protein by pAHD1 or pGPD (Hocher et al 2018) is indicated. Error bars indicate survival standard error (SEM) of at least three independent experiments. Data information: Significance was determined using 2-tailed, unpaired Student’s t test. *P-value 0.01 to 0.05, significant; **P-value 0.001 to 0.01, very significant; ***P-value 0.0001 to 0.001, extremely significant; ****P < 0.0001, extremely significant; P ≥ 0.05, not significant (ns).

Sir3 overexpression led to a 25-fold increase in survival after DSB induction at *TEL6R* which was mainly Dnl4-dependent (Fig 1B). DNA sequencing of repair junctions confirmed that Sir3 overexpression led to increased NHEJ in subtelomeres (Fig 1B and EV1). This effect partly relied on heterochromatin formation since in the absence of Sir4, the NHEJ increase caused by Sir3 overexpression was less pronounced (Fig 1B). However, NHEJ levels in *sir4Δ* cells overexpressing Sir3 remained 7-fold higher than in WT cells, suggesting that Sir3 overexpression also increased NHEJ independently of heterochromatin formation. Consistently, Sir3 overexpression increased NHEJ levels at a euchromatic DSB, although to a more modest extent (Fig 1C, EV1). This data suggests that heterochromatin favours NHEJ repair, and that an excess of Sir3 also stimulates NHEJ in *trans* independently of heterochromatin assembly.

### Sir3 overexpression inhibits MRX-Sae2

Increased NHEJ is a typical phenotype of impaired Mre11 nuclease activity as observed in the *mre11-H125N* nuclease deficient mutant or in absence of its regulator Sae2 (Lee and Lee 2007; Huertas et al. 2008; Huertas and Jackson 2009). Consistently, the absence of *SAE2* and the *mre11-H125N* point mutation led to an epistatic 8-fold increase in NHEJ at euchromatic *TEL6R* and *LYS2* DSB sites (Fig 1B and 1C). We thus tested if Sir3- mediated heterochromatin and the *trans* effect of Sir3 overexpression on NHEJ could result from a defect in Mre11 nuclease activity.

At heterochromatic DSB sites, the deletion of *SAE2* in cells overexpressing Sir3 did not further increase NHEJ, suggesting that MRX-Sae2 is inhibited (Fig 1B). However, *SAE2* deficiency by itself had a significantly lower effect than Sir3 overexpression, indicating that heterochromatin favours NHEJ beyond MRX-Sae2 inhibition. At the euchromatic *LYS2* site, *SAE2* deletion or *mre11-H125N* mutation increased NHEJ in an epistatic manner and to the same extent as Sir3 overexpression (Fig 1C). NHEJ frequencies were not further increased in *sae2Δ* cells overexpressing Sir3 suggesting that Sae2 and Mre11 nuclease activity are inhibited in these cells. Altogether, these results argue that heterochromatin favours NHEJ repair and that the overexpression of Sir3 inhibits MRX-Sae2 in *trans*.

The MRX-Sae2 complex is important to initiate resection of DSB ends (Mimitou and Symington 2008; Garcia et al. 2011; Cannavo and Cejka 2014). To confirm the *trans* inhibition of MRX-Sae2, we tested if Sir3 overexpression could delay resection at a euchromatic site. To assess DSB resection, we employed a PCR-based method to evaluate the resection kinetics at 0.2 and 1 kb kb from the I-SceI cutting site (Fig 1D and EV1; (Batté et al. 2017)). Sir3 overexpression delayed resection of the euchromatic DSB after galactose addition, mimicking the resection delay observed in *sae2Δ* cells (Fig 1E and EV1). The resection delays conferred by *SAE2* deletion and Sir3 overexpression were epistatic (Fig 1E and EV1), consistent with an inhibition of the nuclease activity of the MRX-Sae2 complex upon Sir3 overexpression.

Increased persistence of Mre11 at DSB is typically observed when Mre11 nuclease activity is altered, as seen in *mre11*-*H125N* mutant or in *SAE2* deficient cells (Lisby and Rothstein 2004; Clerici et al. 2006; Cannavo and Cejka 2014; Yu et al. 2018). In agreement with an inhibition of MRX-Sae2 by Sir3, cells overexpressing Sir3 accumulated Mre11 foci following DSB induction (Fig 1F, 1G). The increase in Mre11 foci was comparable to that observed in *sae2Δ* mutants and was not increased upon additional Sir3 overexpression (Fig 1G). Thus, overexpression of Sir3 affects Mre11 turnover at euchromatic DSB sites, recapitulating another typical phenotype of impaired Mre11 nuclease activity. To conclude, Sir3 overexpression increases NHEJ at subtelomeric DSBs through at least two pathways. One that relies on its ability to assemble heterochromatin, and another that limits MRX-Sae2 activity but is independent of heterochromatin formation.

### Sir3 inhibits Sae2 in a dose dependent manner

To dissect the mechanism underlying the inhibition of MRX-Sae2 following Sir3 overexpression, we tested whether this effect was modulated by Sir3 dosage. Under the control of the strong *pGPD* promoter, Sir3 expression increases 29-fold compared to WT. In contrast, under the weaker *pADH1* promoter, Sir3 expression increases only 9-fold (Hocher et al. 2018). We observed that lower Sir3 overexpression resulted in a lesser increase in NHEJ, indicating that Sir3 overexpression impacts NHEJ in a dose dependent manner (Fig 1H).

Upon Sir3 overexpression, Mre11 recruitment to DSB was maintained (Fig 1F), but resection was delayed (Fig 1E), suggesting that Sae2, rather than Mre11, might be the target of Sir3. If true, Sir3 dependent NHEJ increase should be suppressed by Sae2 co-overexpression. To perform Sae2 overexpression, we transformed cells with a high-copy number plasmid bearing the *SAE2* gene under the control of its own promoter. Sae2 overexpression lowers NHEJ levels in *sae2Δ* cells, showing that overexpressed-Sae2 is functional (Fig 1H). Sae2 overexpression partially suppressed the effect of very high Sir3 levels (*pGPD* promoter, 2µ *SAE2*) and completely suppressed the effect of moderately high Sir3 levels (*pADH1* promoter, 2µ SAE2) (Fig 1H). Thus, increased Sae2 expression counteracts the effects of Sir3 overexpression on NHEJ, indicating that Sir3 regulates Sae2 levels or activity.

### Sae2 and Sir3 interact *in vivo* and *in vitro*

Since Sae2 is limiting for normal resection rate (Robert et al. 2011; Tsabar et al. 2015), we addressed the possibility that Sir3 overexpression could regulate cellular levels of Sae2. To do so, GFP fused *SAE2* protein levels were quantified by Western blot in WT or Sir3 overexpressing cells. We observed no major difference in Sae2 protein levels in Sir3 overexpressing cells compared to WT (Fig EV2), indicating that Sir3 overexpression did not impact Sae2 levels.

The dose dependent effect of Sir3 on NHEJ, and its suppression upon increasing Sae2 expression, raises the possibility that Sir3 and Sae2 interact. Consistent with this hypothesis, Sir3 overexpression drastically modified the nuclear distribution of Sae2-GFP. Whereas Sae2-GFP exhibited a diffused nuclear signal in WT cells (Fig 2A), it accumulated in a single bright focus upon Sir3 overexpression (Fig 2A). This bright focus resembled the focus formed by telomeres, Rap1 and SIR proteins in response to Sir3 overexpression (Ruault et al. 2011). Analysis of the localisation of Sae2-GFP and Sir3-mCherry confirmed that the two proteins colocalize in a single cluster upon Sir3 overexpression (Fig 2A), suggesting that they physically interact even in the absence of DSB.

**Figure 2:**
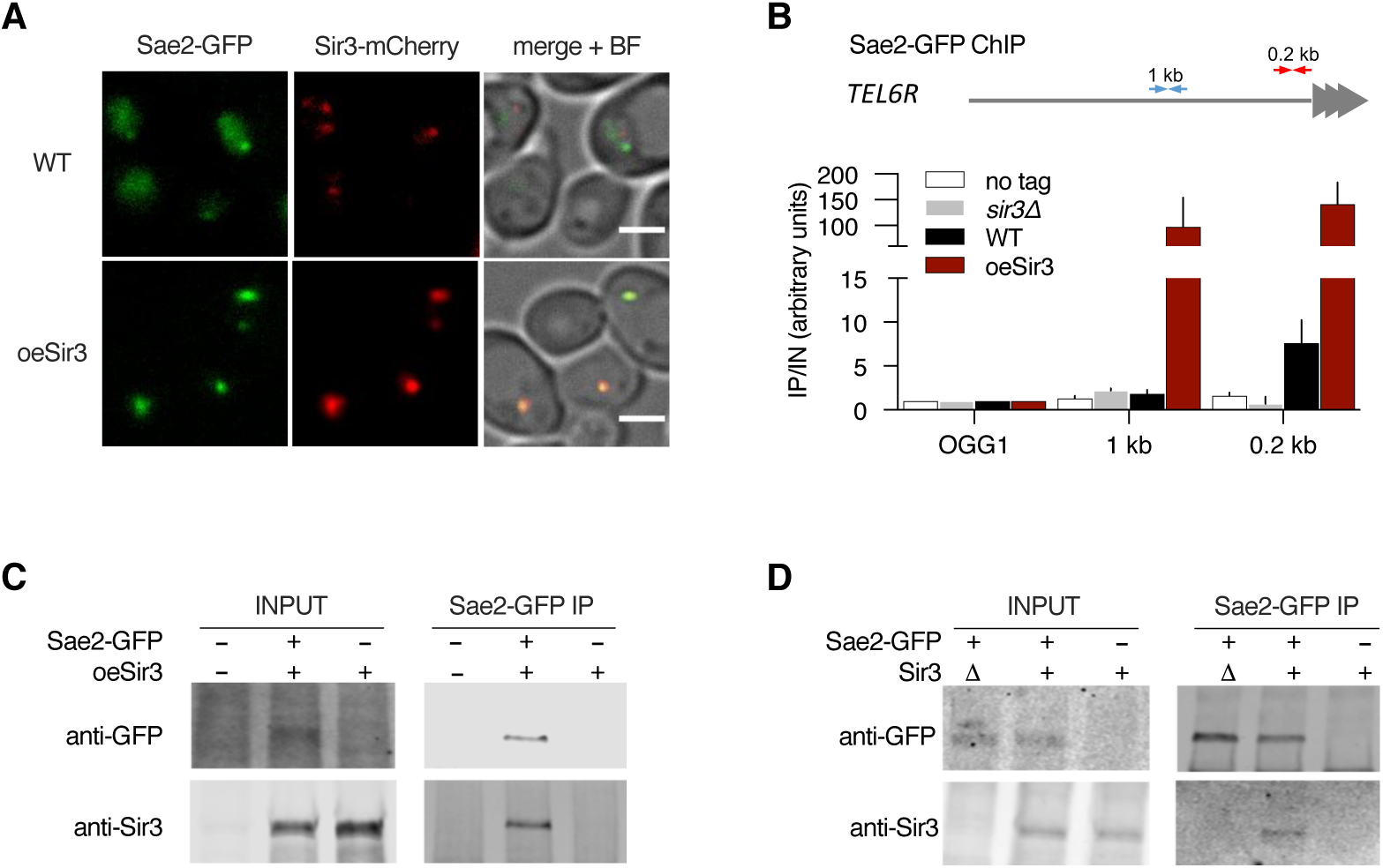
**Sir3 and Sae2 physically interact** A. Representative images of Sir3-mCherry and Sae2-GFP signal in WT and SIR3 overexpressing cells. Scale bars are 2 μm B. Sir3-binding at TEL6R in untagged, WT, sir3Δ cells or in cells overexpressing Sir3 (oeSir3). Binding is probed by ChIP-qPCR 0.2 (red arrows) and 1kb (blue arrows) from telomeres and at the OGG1 control locus using antibodies against Sae2-GFP. The mean of three independent biological replicates is shown and error bars correspond to the variation between replicates. C. Co-immunoprecipitation between Sir3 and Sae2-GFP from cells overexpressing Sir3, analysed by Western blot with anti-GFP and anti-Sir3 antibodies. D. Co-immunoprecipitation between Sir3 and Sae2-GFP from WT cells using antibodies against Sae2-GFP, analysed by Western blot.

Using a chromatin immunoprecipitation (ChIP) approach, we observed Sae2 bound to chromosome ends in WT cells but not in cells lacking Sir3 (Fig 2B). Overexpression of Sir3 increased Sae2 interaction with telomeres and its spreading along subtelomeres suggesting that Sir3 interacts with Sae2 on heterochromatin (Fig 2B).

Sae2-GFP pull-down of Sir3 was achieved in Sir3 overexpressing cells (Fig 2C), and to a lesser extent in WT cells (Fig 2D). This further supports a physical interaction between the two proteins. Furthermore, we observed Sae2-Sir3 interaction using a yeast two-hybrid assay (Fig 3B, 3C and Appendix Fig S1), in agreement with a previous genome-wide screen (Yu et al. 2008), and providing further evidence that the two proteins physically interact *in vivo*. To characterize the domains involved in this interaction, we analysed the multiple sequence alignments of both Sae2 and Sir3 proteins of the *Saccharomycetaceae* family and delineated conserved subdomains (Fig 3A-C, EV3 and Appendix Fig S1). Yeast two-hybrid assays screening of conserved subdomains revealed an interaction between the N-terminal part of Sir3 AAA+ domain (Sir3^SaID^; residues 531-723) and the Sae2 C-terminal domain (Sae2^C^, residues 173-345) (Fig 3A-C, EV3 and Appendix Fig S1). Sir3^SaID^ (for Sae2 Interaction Domain), overlaps with the previously defined Sir4 interacting domain (Fig 3A; (King et al. 2006)). However, Sir4 was not required for the observed Sir3-Sae2 two-hybrid interaction (Fig 3D).

**Figure 3:**
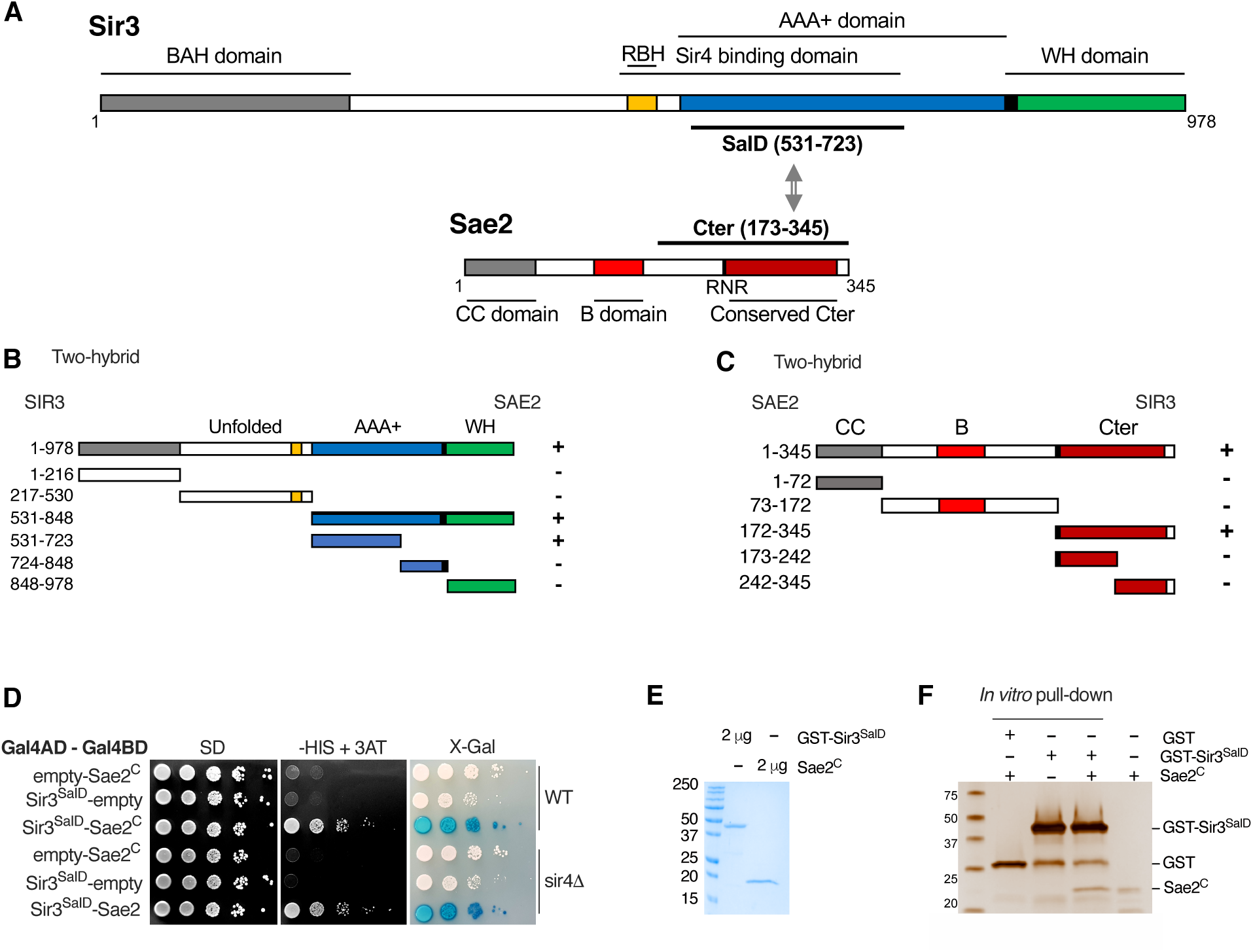
**Direct physical interaction between Sir3^SaID^ and Sae2^C^ domains** A. Schematic representation of Sir3 and Sae2 protein domains. B. Delineation of the Sir3 domain responsible for interaction with Sae2 by two-hybrid assays. The GAL4-BD fusions with indicated Sir3 fragments were tested in combination with a GAL4–AD–Sae2 fusion; “+” indicates an interaction. C. Delineation of the Sae2 domain responsible for interaction with Sir3 by two-hybrid assays. The GAL4-BD fusions with indicated Sae2 fragments were tested in combination with a GAL4–AD–Sir3 fusion; “+” indicates an interaction. D. Yeast two-hybrid interaction analysis between Sae2^C^ and Sir3^SaID^ domains in WT or sir4Δ cells. Growth on -His + 3AT and blue coloration on X-gal indicate an interaction. E. Comassie-stained gel of purified GST-Sir3^SaID^ and Sae2^C^ peptides. F. Representative silver-stained gels of in vitro GST-pulldown of GST or GST-Sir3^SaID^ and Sae2^C^ purified peptides. Control: Sae2^C^ (300 ng, lane 4).

To verify that the Sir3-Sae2 interaction was direct, we purified histidine-tagged Sae2^C^ and GST-tagged Sir3^SaID^ fragments expressed in bacteria (Fig 3E) and performed *in vitro* pull-down experiments. Sae2^C^ was retrieved with purified GST-Sir3^SaID^, but not with GST alone showing specific direct interaction (Fig 3F). Protein extracts used for this experiment were supplemented with benzonase to remove DNA, showing that DNA did not mediate the interaction and that direct protein interaction takes place between Sir3^SaID^ and Sae2^C^. Altogether, these results show that Sae2 directly interacts with Sir3. This interaction might be the basis of the Sae2 inhibition observed upon Sir3 overexpression.

### Sae2 and Sir3 interaction prevents Sae2 functions and promotes NHEJ

To functionally test whether Sae2-Sir3 interaction inhibits Sae2, we screened for Sir3 mutants without the capacity to interact with Sae2. To this end, we designed a two-hybrid screen to select separation of function Sir3 mutants no longer interacting with Sae2, while retaining interaction with Sir4. For this, we used a strain in which *HIS3* and *LacZ* reporter genes associate with *GAL1* UAS and *lexAop* DNA targeting sequences respectively. This allows for the simultaneous assessment of a positive interaction between two proteins, alongside the loss of interaction between one of those proteins and a third (Fig 4A). We transformed this strain with plasmids expressing *SIR4^C^* fused to a *GAL4* binding domain and *SAE2^C^* fused to a *LexA* binding domain that bind upstream of *HIS3* and *LacZ* respectively. A Sir4 binding partner fused to the Gal4 activating domain will thus activate the expression of *HIS3*, whereas a Sae2 binding partner fused to the Gal4 activating domain will activate the expression of *LacZ*.

**Figure 4:**
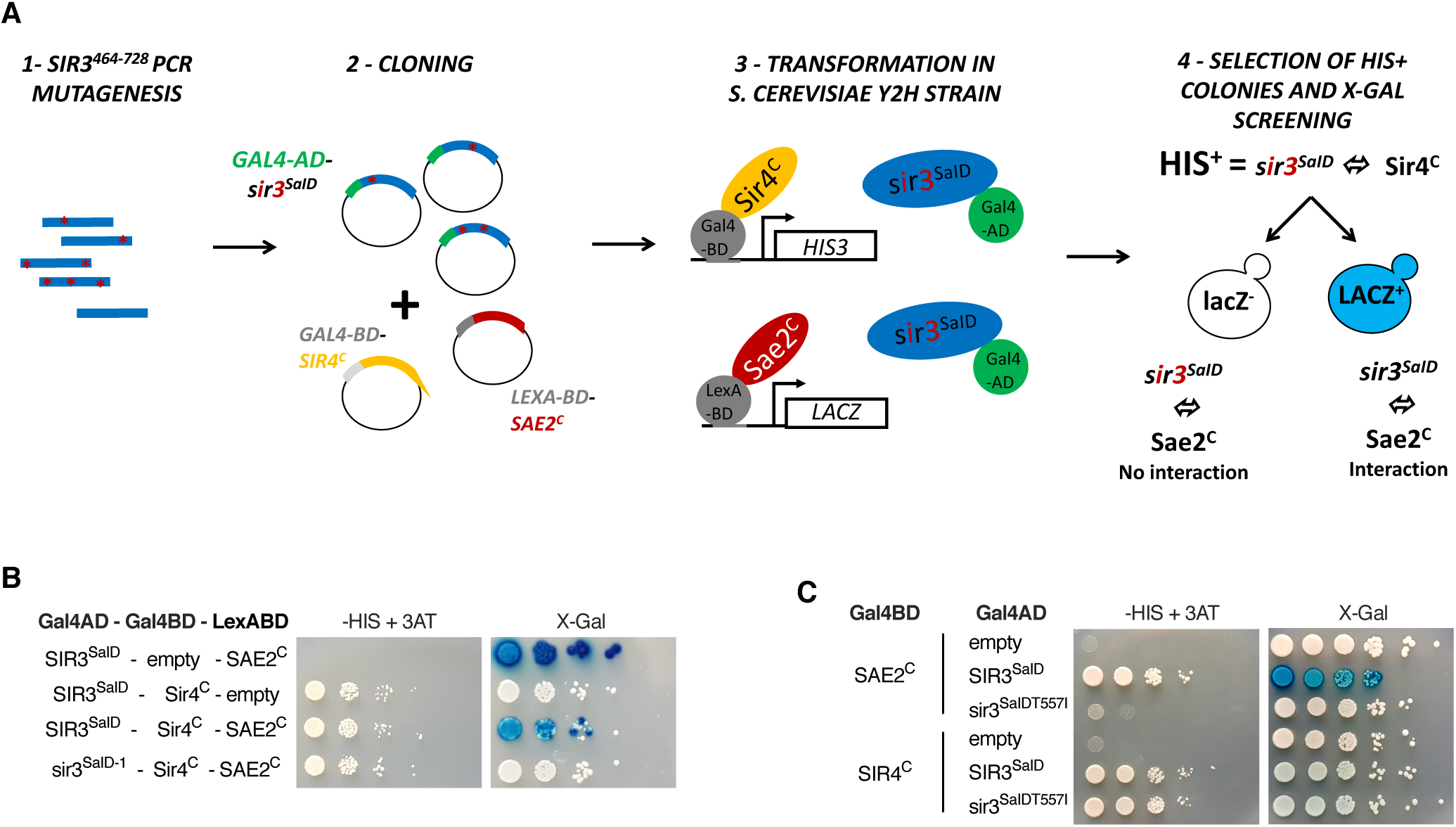
**The T557I point mutation in Sir3 abolishes Sae2-Sir3 interaction** A. Schematic representation of the assay used to screen for SIR3 mutants deficient for Sae2 interaction while maintaining interaction with Sir4. The SIR3^SaID^ fragment (464-728) was mutagenized by PCR, cloned in the pACT2 two hybrid plasmid and transformed into the reporter strain along with plasmids expressing LexA-BD-SAE2^C^ and GAL4-BD-SIR4^C^ fusion proteins. The reporter strain (yKD1991) bears a Gal4 binding sequence (Gal4BD) upstream of a HIS3 reporter gene, and a LexA binding sequence (LexABD) precedes a LacZ reporter gene. Transformants in which the gal4-BD-Sir4^C^ and Sir3^SaID^-gal4-AD fragments interact were selected for HIS3 expression on -HIS + 3AT medium and subsequently screened for LacZ expression upon X-gal coloration. Cells showing no LacZ expression were collected and the mutated sir3^SaID^-GAL4-AD was retrieved and sequenced. B. Representative images of two hybrid assays in the yKD1991 strain testing the interaction of the WT or the mutant SIR3^SaID^ fragment isolated from the screen with SAE2^C^ or SIR4^C^. C. Representative images of two hybrid assays testing the interaction of the WT or the mutant SIR3^SaIDT557I^ fragment with SAE2^C^ or SIR4^C^.

We performed random mutagenesis of a Sir3 domain sufficient to interact with Sae2 and Sir4 (464-728 aa;(King et al. 2006)), and created a library of mutated *SIR3^SaID^* fused to the *GAL4* activating domain (GAD). This library was introduced into the screening strain and Sir3 mutants still able to interact with Sir4 were selected based on their ability to grow on media lacking histidine and supplemented with aminotriazole (-HIS + 3-AT). This step eliminates non-sense or non-expressed GAD-SIR3^SaID^ mutants. X-Gal staining of His+ colonies allowed for the selection of white clones in which the GAD-SIR3^SaID^ - LexABD- Sae2^C^ interaction was lost.

Using this screen, we recovered a mutant deficient for Sir3-Sae2 interaction while proficient for Sir3-Sir4 interaction. Sequencing of this mutant identified two point mutations T557I and T598A followed by a frameshift at position 707 (sir3^SaID-1^; Fig 4B). These two residues are not strictly conserved among the *Saccharomycetaceae* family, but T557 is flanked by a conserved patch (Fig EV3). Subcloning of the individual mutations and secondary two-hybrid tests showed that the mutation T557I alone is sufficient to impair the Sae2-Sir3 interaction while preserving the Sir4-Sir3 interaction (sir3^SaID-T557I^; Fig 4C).

To test the functional consequences of Sir3-Sae2 interaction loss, we assessed NHEJ in strains overexpressing either the WT or T557I mutant Sir3^SaID^ fragment (Fig 5A). High-level expression of the Sir3^SaID^ fragment was sufficient to promote NHEJ and displayed an epistatic relationship with the loss of Sae2 (Fig 5A and EV4). In contrast, high-level expression of the mutated fragment had no effect, indicating that Sae2 inhibition by Sir3^SaID^ requires an intact Sae2-Sir3 interaction (Fig 5A).

**Figure 5:**
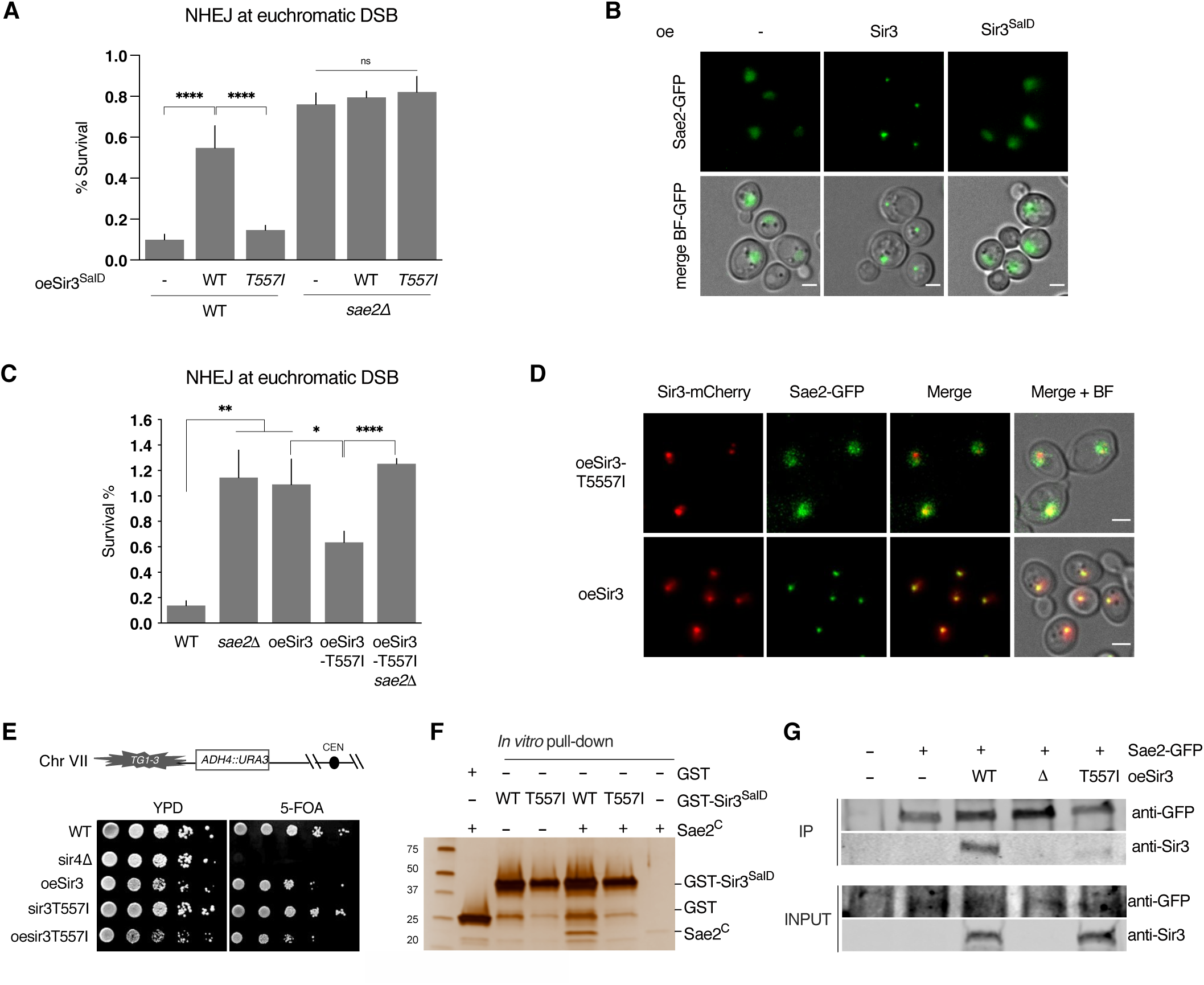
**Sir3-Sae2 interaction prevents Sae2 function and promotes NHEJ** A. Survival frequencies after DSB induction at LYS2 locus in WT or sae2Δ strains where the Sir3^SaID^ or Sir3^SaIDT557I^ domains are overexpressed from a GPD promoter at the SIR3 locus. Error bars indicate survival standard error (SEM) of at least three independent experiments. B. Representative images of Sae2-GFP in WT cells and in cells overexpressing either full-length Sir3 or the sir3SaID domain. Scale bars are 2 μm C. Survival frequencies after DSB induction at LYS2 locus in the indicated strains. Error bars indicate survival standard error (SEM) of at least three independent experiments. D. Representative images of Sir3-mCherry and Sae2-GFP signal in cells overexpressing Sir3 and sir3T557I . Scale bars are 2 μm. E. Telomeric silencing assay at TEL7L in WT, sir4Δ, sir3T557I cells, cells overexpressing SIR3 (oeSIR3) or sir3T557I (oesir3T557I). Growth on 5-FOA plates reflects telomeric silencing. F. Representative silver-stained gels of in vitro GST-pulldown of GST or GST-Sir3^SaID^, GST-sir3-T557I^SaID^ and Sae2^C^ purified peptides. Control: Sae2^C^ (300 ng, lane 6). G. Co-immunoprecipitation between Sae2-GFP and Sir3 from untagged, Sae2-GFP WT cells, and Sae2-GFP cells overexpressing WT Sir3 (oeSir3, WT), Sae2-GFP sir3Δ or Sae2- GFP overexpressing the sir3-T557I mutant (oeSir3, T557I) using antibodies against Sae2- GFP, analysed by Western blot with anti-GFP and anti-Sir3 antibodies. Data information: Significance was determined using 2-tailed, unpaired Student’s t test. *P-value 0.01 to 0.05, significant; **P-value 0.001 to 0.01, very significant; ***P-value 0.0001 to 0.001, extremely significant; ****P < 0.0001, extremely significant; P ≥ 0.05, not significant (ns).

Strikingly, overexpression of the Sir3^SaID^ fragment, which is sufficient to inhibit Sae2 (Fig 5A), did not promote Sae2 clustering (Fig 5B). This shows that Sae2 inhibition by Sir3 is maintained, even when Sae2 is not trapped in the telomere cluster. These results indicate that the inhibition of Sae2 is not only a secondary consequence of its sequestration by Sir3, but rather suggests that Sir3-Sae2 interaction *per se* can inactivate Sae2.

Insertion of the *T557I* mutation in the full-length *SIR3* gene reduced the ability of Sir3 to promote NHEJ when overexpressed (Fig 5C). This correlated with a loss of Sae2-sir3- T557I colocalization (Fig 5D). Importantly, the point mutation does not affect the stability of Sir3, and sir3-T557I overexpressing cells retained the ability to form the telomere hypercluster (Fig 5D and EV4) and propagate subtelomeric heterochromatin (Fig 5E). In contrast, Sae2-GFP no longer formed a single bright focus in sir3-T557I overexpressing cells (Fig 5D), indicating that Sae2 clustering requires Sae2-Sir3 interaction. *In vitro* the purified Sir3^SaiD^ mutated fragment failed to interact with Sae2^C^ (Fig 5F). However, the residual NHEJ observed in cells overexpressing full length sir3-T557I suggests that it retains some interaction with Sae2 *in vivo* (Fig 5C). This was confirmed by co-immunoprecipitation experiments that showed a residual interaction with Sae2 *in vivo* (20±10 % of the interaction detected in WT, Fig 5F). Altogether, these data show that the Sir3^SaID^ domain is sufficient to interact with and inhibit Sae2, and that interaction between Sir3 and Sae2 is necessary and sufficient to inhibit Sae2 activity.

### Sae2 and Sir4 compete for Sir3 binding

To explore further the functional consequences of the Sir3-Sae2 interaction, we assessed NHEJ in the absence of the SIR complex. Strains used lack the *HML* locus to avoid indirect effects on NHEJ efficiency caused by pseudo-diploidization, as observed following the derepression of the cryptic mating type loci in strains with *SIR* deletions (Aström et al. 1999; Lee et al. 1999; Frank-Vaillant and Marcand 2001). Consistent with an inhibition of Sae2 by Sir3 expressed at physiological levels, NHEJ was reproducibly decreased by ∼2- fold in *sir3Δ* mutants both at chromosomic DSB sites and in plasmid rejoining assay (Fig 6A and 6B). In contrast, *sir4Δ* mutants exhibited a more than 2-fold increase in NHEJ relative to WT, which was abolished by the additional loss of Sir3 (Fig 6A). This increase was epistatic with *sae2Δ*, suggesting that Sae2 and Sir4 act in the same pathway to inhibit NHEJ. Together, these results show that Sir3 is required to increase NHEJ in absence of Sir4 and suggest that the regulation of Sae2 by Sir3 is involved. Consistently, the sir3-T557I mutant impaired for interaction with and inhibition of Sae2, fails to increase NHEJ in absence of Sir4 (Fig 6A). Therefore, physiological levels of Sir3 are sufficient to inhibit Sae2, and Sir4 is able to counteract this inhibition.

**Figure 6:**
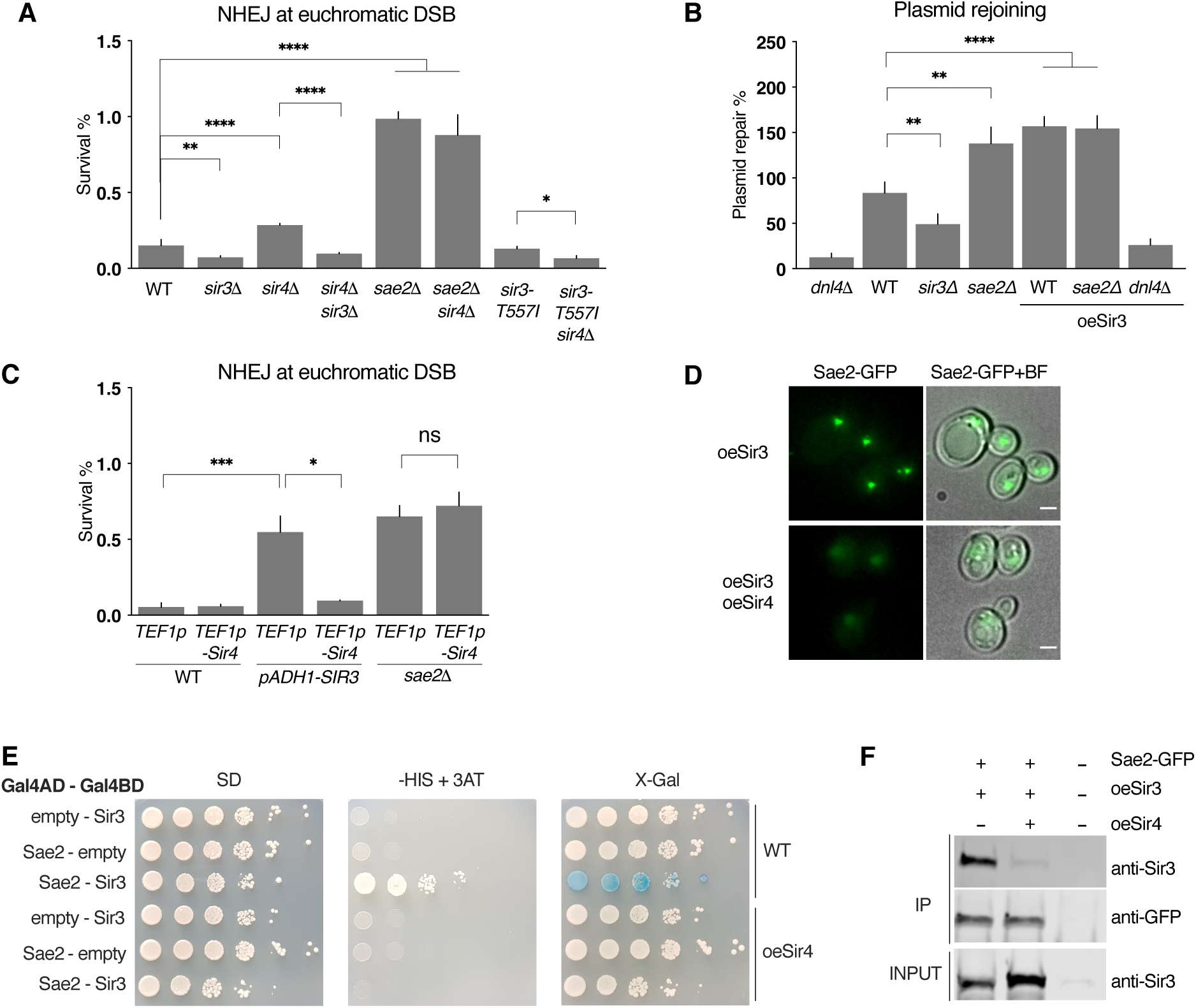
**Sir3-Sae2 interaction is modulated by Sir4.** A. Survival frequencies after DSB induction at LYS2 locus in the indicated strains. Error bars indicate survival standard error (SEM) of at least three independent experiments. B. Plasmid cleaved by Xho I was transformed into strains and NHEJ efficiency was measured. Error bars indicate survival standard error (SEM) of at least three independent experiments. C. Survival frequencies after DSB induction at LYS2 locus in the indicated strains. Insertion of the strong TEF1p promoter upstream of the SIR4 ORF leads to Sir4 overexpression. Insertion of the ADH1p promoter upstream of SIR3 leads to mild Sir3 overexpression. Error bars indicate survival standard error (SEM) of at least three independent experiments. D. Representative images of Sae2-GFP in cells overexpressing Sir3 (oeSir3) and expressing or not high levels of Sir4 (oeSir4). Scale bars are 2μm E. Representative images of two hybrid assays testing the interaction between the full-length Sir3 and full-length Sae2 proteins in WT cells expressing or not high levels of Sir4 (oeSir4 and WT respectively). F. Co-immunoprecipitation between Sae2-GFP and Sir3 from untagged, Sae2-GFP cells overexpressing Sir3 and expressing or not high levels of Sir4 (oeSir4). Data information: Significance was determined using 2-tailed, unpaired Student’s t test. *P-value 0.01 to 0.05, significant; **P-value 0.001 to 0.01, very significant; ***P-value 0.0001 to 0.001, extremely significant; ****P < 0.0001, extremely significant; P ≥ 0.05, not significant (ns).

As Sae2 and Sir4 interact with the same Sir3 domain (Fig 3A), a competition between Sir4 and Sae2 for Sir3 binding might explain NHEJ increase in cells lacking Sir4 and the dependence of this increase on Sir3 and Sae2. If Sir4 and Sae2 compete for Sir3 binding, overexpression of Sir4 should prevent Sir3-Sae2 interaction and counteract the increase in NHEJ caused by Sir3 overexpression. We tested this hypothesis by Sir4 overexpression, through genomic insertion of an additional copy of the *SIR4* gene under the control of a strong promoter (*TEF1p*). As predicted, Sir4 overexpression alongside Sir3 overexpression restored NHEJ to WT levels whereas it did not affect NHEJ in WT or *sae2Δ* cells (Fig 6C). This indicates that Sir4 overexpression does not affect NHEJ by itself, but instead counteracts Sir3-overexpression-mediated inhibition of Sae2. Expressing high levels of Sir4 was also sufficient to counteract the Sir3-overexpression induced clustering of Sae2 (Fig 6D), to disrupt the two-hybrid interaction detected between Sir3 and Sae2 (Fig 6E) and to decrease co-immunoprecipitation of Sir3 with Sae2 (Fig 6F), showing that Sir4 binding to Sir3 counteracts Sir3-Sae2 interaction. Note that the favoured partner of Sir3 at subtelomeres remains Sir4, since Sae2 overexpression had no effect on silencing (Fig EV5). Collectively, this data is consistent with a model in which Sae2 is inactive when bound to Sir3, but can be released by the competitive binding of Sir4 to Sir3.

### Sir3-mediated Sae2 inhibition mechanism

How might Sir3-Sae2 interaction inhibit Sae2? Interestingly, the C-terminus of Sae2, which we have demonstrated as sufficient for Sir3 interaction, also interacts with Rad50. This interaction requires Sae2 C-terminus phosphorylation and is essential for stimulation of Mre11 nuclease activity (Cannavo and Cejka 2014; Cannavo et al. 2018). Sir3 binding to Sae2 could thus impair the interaction between Sae2 and MRX^MRN^ by steric hindrance, or by impairing Sae2 C-terminus phosphorylation.

If Sir3 impairs Sae2-MRX^MRN^ interaction, Sae2 recruitment to DSB, which relies on its interaction with MRX (Mojumdar et al. 2021), should be affected. Thus, we monitored Sae2 association with DSB by chromatin immunoprecipitation (ChIP) upon Sir3 overexpression. We observed lower enrichment of Sae2 at DSB in Sir3 overexpressing cells compared to WT (Fig 7A) while Sae2 binding to telomeres was not significantly affected (Fig 7B). These data support the hypothesis that the trapping of Sae2 in the Sir3-mediated telomere cluster impairs its ability to interact with MRX and to be recruited at DSB sites. In contrast, overexpression of the Sir3^SaID^ domain which is sufficient to inhibit Sae2 (Fig 5A) did not sequester Sir3 at telomeres (Fig 7B) and lead to a greater enrichment of Sae2 at DSB (Fig 7A). These results suggest that Sae2-MRX interaction is not affected by Sir3^SaID^-Sae2 interaction *per se* when Sae2 is not trapped in the telomere cluster, and support the hypothesis that Sir3-Sae2 interaction is sufficient to inhibit Sae2 activity. In agreement with Sae2 inhibition by Sir3, independently of Sae2 binding to MRX at DSB, the absence of Sir3 does not significantly affect Sae2 enrichment at DSB (Fig 7A).

**Figure 7:**
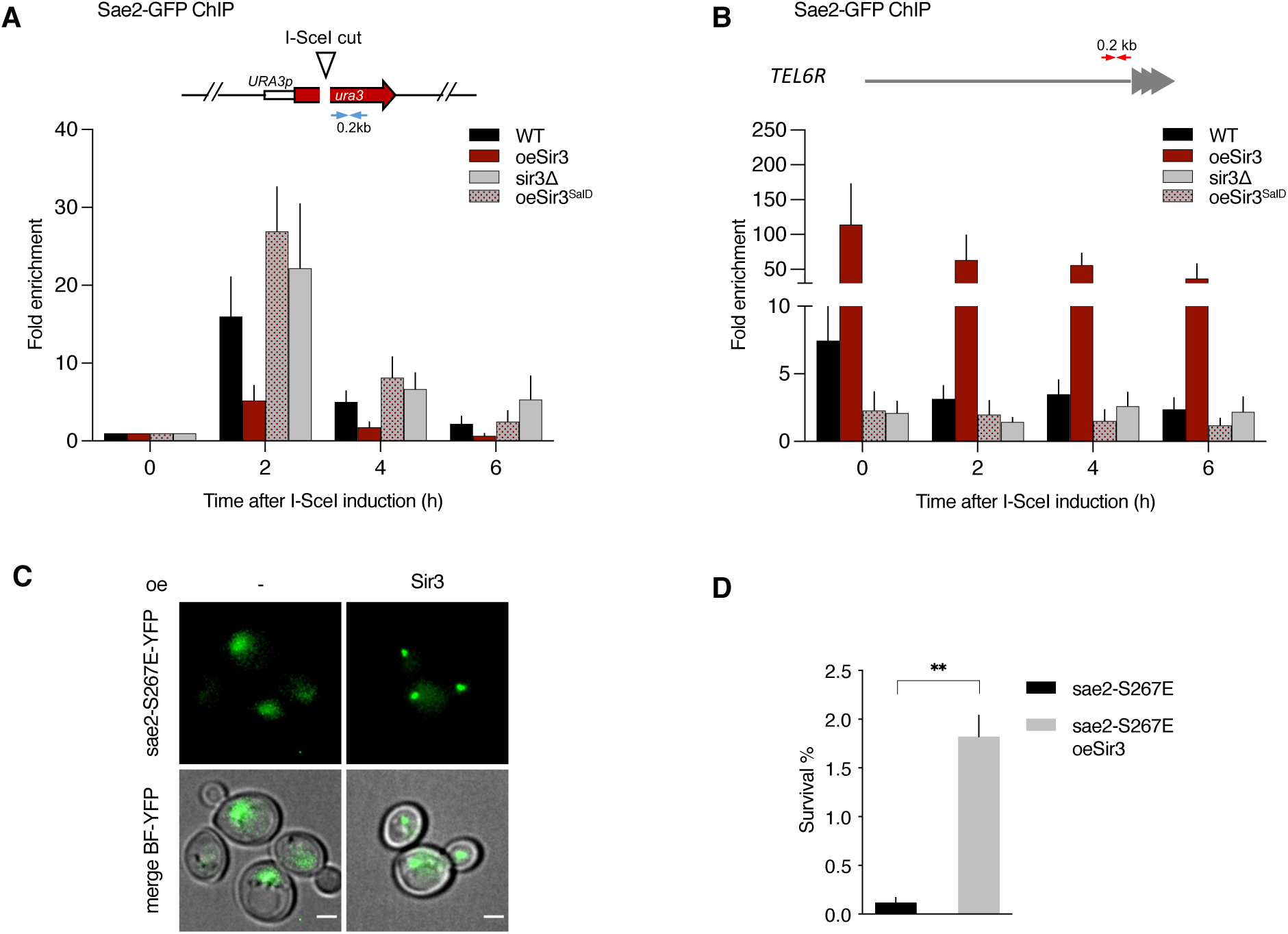
**Modulation of Sae2 recruitment and activity by Sir3** A. Relative fold enrichment of Sae2 at 0.2 kb from I-SceI site was evaluated by qPCR after ChIP with anti-GFP antibodies. The error bars indicate the variation between at least three biological replicas. B. Relative fold enrichment of Sae2 at 0.2 kb from *TEL6R* after DSB induction at the LYS2 I-SceI site was evaluated by qPCR after ChIP with anti-GFP antibodies. The error bars indicate the variation between at least three biological replicas. C. Representative images of sae2-E267E mutant cells overexpressing or not full-length Sir3. Scale bars are 2 μm D. Survival frequencies after DSB induction at LYS2 locus in *sae2-E267E* mutant cells overexpressing or not full-length Sir3 (oeSir3). Data information: Significance was determined using 2-tailed, unpaired Student’s t test. *P-value 0.01 to 0.05, significant; **P-value 0.001 to 0.01, very significant; ***P-value 0.0001 to 0.001, extremely significant; ****P < 0.0001, extremely significant; P ≥ 0.05, not significant (ns).

As stated above, Sir3 could inhibit Sae2 activity by preventing the phosphorylation of the conserved CDK site at S267 of Sae2 C-terminus which is critical for its resection function *in vitro* and *in vivo* (Huertas et al. 2008; Cannavo and Cejka 2014; Cannavo et al. 2018; Zdravković et al. 2021). If this were the case, Sir3 inhibition should be relieved by the *sae2-S267E* phosphomimic mutant. However, in contrast to this prediction, sae2-S267E interacts with overexpressed Sir3 in the telomere cluster (Fig 7C) and is inhibited by Sir3 overexpression (Fig 7D). Altogether, these results indicate that Sir3 inhibits Sae2 once it has been phosphorylated by CDK and is bound to MRX, and suggest that Sir3 affects the subsequent step of mre11 nuclease activation.

## Discussion

Initiation of DSB end resection is a pivotal decision during DSB repair: it precludes the “by default” repair by NHEJ and commits cells to repair by HR (Frank-Vaillant and Marcand 2002; Symington 2016). The effector of this irreversible pathway choice decision, the MRX-Sae2 complex, is the focus of various regulatory inputs, including cell cycle phase (Huertas et al. 2008; Huertas and Jackson 2009; Cannavo and Cejka 2014). Here we reveal an unexpected role for Sir3 in impinging on this pathway choice. Sir3 physically interacts with Sae2, inhibits its DSB-end resection initiation function and consequently increases MRX retention at DSB sites and NHEJ efficiency.

Altogether, our results suggest that Sae2 inhibition relies on the inactivation of Sae2 following Sir3 binding to its C-terminus, the efficiency of which depends on the relative abundance of each protein. Sir3 binding *per se* does not inhibit Sae2 recruitment to DSB and hence Sae2 interaction with MRX, but counteracts the ability of Sae2 to stimulate MRX nuclease activity. Sae2 has been shown to bind MRX^MRN^ through several independent interactions involving Mre11, Xrs2 and Rad50 (Cannavo and Cejka 2014; Liang et al. 2015; Cannavo et al. 2018). Sir3 is unlikely to affect Sae2 interaction with Mre11, which fails to interact with Sae2 C-terminal fragment *in vitro* (Cannavo et al. 2018), or with Xrs2, which interacts with Sae2 N-terminus (Liang et al. 2015). It could however play on the binding of Sae2 C-terminus to Rad50 heads, which is sufficient to stimulate Mre11 nuclease activity (Cannavo et al. 2018).This hypothesis is further supported by the phenotype of the *rad50S* mutant, which is reminiscent of the phenotype caused by Sir3-Sae2 interaction. Indeed, the *rad50S*(K81I) mutation which affects Rad50-Sae2C interaction *in vitro* (Cannavo et al. 2018) impairs Sae2-mediated MRX activation, but still allows strong recruitment of Sae2 to DSB *in vivo* (Yu et al. 2018). Based on these observations, a hypothesis that awaits closer examination is that Sir3 binding to Sae2 could impair the Sae2-Rad50 interaction required for Sae2-mediated stimulation of Mre11 endonuclease activity.

In addition, our data suggest that a pool of Sir3-bound Sae2 in subtelomeric chromatin is prevented from activating MRX nuclease activity. One rationale behind limiting the availability of Sae2 could be to limit resection, considering NHEJ is sufficient for the repair of most DSB. Uncontrolled resection might also drive repair towards error-prone HR (SSA, BIR) leading to loss of genetic information (Chen et al. 2013; Toledo et al. 2013; Lee et al. 2016; Batté et al. 2017). Thus, tight control of the Sae2 pool that can engage in end processing is needed to ensure genome integrity. The Sir3-bound pool of Sae2 at subtelomeres is inactive, which limits resection and promotes NHEJ, ensuring genetic integrity of subtelomeres. Sir3 impact at euchromatic DSB sites suggests that release of this pool in response to DSB in regulating this pool could also have a more general role in DNA repair. Consistently, the Sir3-bound pool of Sae2 at subtelomeres decreased in response to DSB induction. Although release of Sae2 from telomeres might have been expected to increase Sae2 DSB binding, the absence of Sir3 does not significantly improve Sae2 recruitment to DSB. This may not be surprising when considering that an excess of Sae2 increases MRX turnover at DSB (Yu et al. 2018) and may in turn limit Sae2 binding.Interestingly, our results echo previous studies showing that the SIR complex dissociates from telomeres in S-G2 phase or upon damage induction (Martin et al. 1999; Mills et al. 1999; McAinsh et al. 1999). Whether Sae2 and Sir3 are released from telomeres as a complex remains to be determined experimentally. One possibility, considering that the SIR complex in solution contains Sir2, Sir3 and Sir4 in a 1:1:1 molar ratio (Cubizolles et al. 2006), is that Sir3 once released from the telomeres is not able to accommodate Sae2 binding. Release of SIRs from telomeres could thus liberate a pool of Sae2 free to act at DSB. A non exclusive scenario is that Sir3 which also associates with DSBs (Martin et al. 1999; Mills et al. 1999) controls Sae2 activity at DSB, limiting MRX activity to prevent excessive resection. A timely regulation of Sae2-Sir3 interaction and of their release from telomeres could participate in the fine regulation of active Sae2 at DSB.

Sae2-Sir3 interaction may also be relevant for telomere length regulation. During telomere replication, Sae2 has a facultative role in facilitating the generation of the G rich 3’- ssDNA, the telomerase substrate, and therefore in promoting telomere elongation (Bonetti et al. 2009). Interestingly, in cells lacking Tel1, where telomerase recruitment depends exclusively on Mec1 (Arnerić and Lingner 2007), and therefore possibly more so on resection, Sae2 loss slightly shortens telomeres, compared to Sir3 loss, which elongates them (Appendix Fig S2). In tel1Δ cells lacking Sae2, Sir3 loss does not impact telomere length, suggesting that the telomere elongation observed in the presence of Sae2 might be a consequence of increased Sae2 activity. Although the network of interactions at telomeres does not allow us to rule out indirect effects, this data suggests that Sir3 could downregulate Sae2 at telomeres.

The inhibition of Sae2 by Sir3 is suppressed by Sir4 overexpression suggesting that Sir4 competes with Sae2 for Sir3 binding, and that this competition modulates the inhibition of Sae2 by Sir3. This competition may explain how Sir4 loss increases NHEJ in euchromatin (Fig 6A), simply by increasing the pool of Sae2 associated with and inhibited by Sir3. This Sir4-dependent Sae2 activation could also promote telomere protection against NHEJ if telomere-associated Sir3 molecules are in complex with Sir4.

The competition between Sae2 and Sir4 for Sir3 binding questions the relevance of Sae2-Sir3 interaction in subtelomeric heterochromatin. Recent *in vitro* data support a stoichiometry of two Sir3 molecules and one Sir2–4 dimer per nucleosome (Swygert et al. 2018). This suggests that one Sir3 molecule per nucleosome might not be interacting with Sir4 on chromatin, leaving room for the binding and inhibition of Sae2 on heterochromatin-bound Sir3. Consistently, we detect a Sir3-dependent Sae2 binding at subtelomeres in WT cells (Fig 2B), and the inhibition of Sae2 activity at heterochromatic subtelomeric DSB (Fig 1A). The main function of Sir3-mediated Sae2 inhibition could thus be to protect subtelomeres from resection, avoiding loss of genetic information and providing chromosome end deprotection.

NHEJ is favoured at heterochromatic DSB, beyond Sae2 inhibition by Sir3, through a mechanism that remains to be defined. The presence of the NHEJ factor KU at subtelomeres (Martin et al. 1999), mediated by its interaction with Sir4 (Roy et al. 2004), could favour NHEJ. Alternatively, the fact that heterochromatin limits resection, likely beyond MRX-Sae2 inhibition, may stabilise unprocessed DSB ends, therefore increasing NHEJ likelihood. It is striking to note that NHEJ is favoured in heterochromatic subtelomeres, despite its strong inhibition at telomere ends (Marcand et al. 2008). This dichotomy is conserved in mammalian cells in which NHEJ is prevented at telomeres, but not near them (van Steensel et al. 1998; Muraki et al. 2015). At yeast telomeres, a key NHEJ repressor is Sir4, which acts, at least in part, in a Sir3 independent manner (Marcand et al. 2008; Roisné-Hamelin et al. 2021; Khayat et al. 2021). Sir4 thus seems to have two opposite functions in NHEJ regulation depending on chromosomic context: a strong repressive function at telomeres, and a stimulating function at subtelomeres. Several hypotheses can be proposed to account for these differences. Sir4 could be present in different amounts at subtelomeres and at telomeres. Alternatively, Sir4 could adopt distinct conformations that would dictate its ability to inhibit NHEJ depending on its binding partners at telomeres compared to subtelomeres.

In mammals, NHEJ is the prevalent repair mechanism in non-coding and silent chromatin (Aymard et al. 2014), and in perinuclear heterochromatin (Lemaître et al. 2014). Furthermore, CtIP interacts with BARD1, a HP1 binding partner, as well as CBX4, an E3 SUMO ligase subunit of the facultative heterochromatin Polycomb complex (Wu et al. 2015; Soria-Bretones et al. 2017). Whether this is associated with regulation of CtIP activity remains to be investigated. This data, together with our observations suggest that a regulation of the MRX^MRN^-Sae2^CtIP^ complex by the chromatin context might be a conserved general principle.

Here, we provide the first insights into the mechanisms regulating DSB repair in yeast heterochromatin. We show that the early resection step, which controls the choice between NHEJ and HR, is tightly regulated in heterochromatin. Notably, there is a stringent regulation of the MRX^MRN^ complexes potent end-resection activity, through the direct inhibition of Sae2^CtIP^ by Sir3. Precise characterization of the Sir3-Sae2^CtIP^ binding interface will help in understanding how Sae2 binds and activates MRX and may enable the design of specific synthetic inhibitors towards Sae2^CtIP^-mediated MRX^MRN^ activation, the cornerstone of HR/NHEJ repair pathway choice.

## Methods

### Plasmids

Two-hybrid plasmids (*pACT2*-*SAE2*, *pACT2-SIR3^SaID^*, *pACT2-SIR3, pACT2-SIR3^464-728^, pGBT9-SIR3, pGBT9-SAE2^C^, pGBT9-SIR3^SaID^, pLexA-SAE2^C^)* were constructed by inserting the full length or appropriate fragments of *SAE2* and *SIR3* genes, amplified from W303 genomic DNA, in pACT2, pGBT9 and pBTM116 vectors digested by BamHI by single strand annealing cloning (SLIC, (Li and Elledge 2007)). To test interactions with Sir4, the pGBD-C2- SIR4 plasmid was used (Ehrentraut et al. 2011). *pACT2-sir3^SaID-T557I^* and *pACT2-sir3-T557I* were generated by rolling circle mutagenesis of *pACT2-SIR3^SaID^* and *pACT2-SIR3* as described in (Hansson et al. 2008). To overexpress Sae2, the *SAE2* gene was amplified from W303 genomic DNA and inserted in pRS423 digested by *SalI*-HF by SLIC (Li and Elledge 2007) to produce pKD343. To overexpress *SIR4* for NHEJ assays*, SIR4* amplified from W303 genomic DNA was inserted by SLIC in pKD431, an integrative plasmid pRS403 with a TEF1p promoter, to generate pKD432. Genomic integration of the plasmids at *HIS3* is possible after digestion by *PstI*.

The *SIR3^SaiD(531-723)^* fragment was cloned under the T7 promoter into the vector pnEAvG (Diebold et al. 2011) generating pKD434 that allows GST- Sir3^SaID^ expression in bacteria. *SAE2^C^* was cloned into an adapted SUMO vector (pKD435) allowing His6-SUMO-Sae2^Cter^ protein expression.

Mutagenesis of the sequence encoding Sir3^464-728^ using the GeneMorph II EZClone Domain Mutagenesis Kit (Agilent, 200552-5) was performed by PCR on 9,5 µg of *pACT2-SIR3^464-728^* with 20 cycles of amplification to allow low mutation rate. The PCR products were subsequently subcloned by SLIC in pACT2 and Sanger sequenced for mutation rate estimation.

### Yeast strains

All strains in this study are isogenic to W303 (Mata (or Matα) ADE2 leu2-3,112 his3-11,15 trp1-1 ura3-1) and are listed in Appendix Table S1. For DSB induction the *I-SCEI* gene was introduced in the yeast genome by transformation of the cells with pKD144 (pRS404-*GAL1p-I-SceI*) digested by *PmlI* to insert in *TRP1*. Gene deletions and insertions of strong constitutive promoters (GPDp, ADH1p) were performed by PCR-based gene targeting (Longtine et al. 1998).

The mre11-H125N allele was introduced in strains by crossing with the LSY2854-21C strain (Chen et al. 2013). Mre11-YFP was introduced in strains by cross with the W5089-6A strain (Kaiser et al. 2011). Sae2-AAGRRIGDGAGLIN-GFP was constructed by PCR gene targeting on pKT128 (Sheff and Thorn 2004). *SIR3*-*mCherry* was constructed by PCR gene targeting on pSL1 (Léon et al. 2008) whose marker was replaced by Hygromycin B resistance (HPH^r^), with primers pr1328 and pr1329.

### Media and growth conditions

Yeast strains were grown in rich medium (yeast extract–peptone–dextrose, YPD) or synthetic complete (SC) medium lacking the appropriate amino acid at 30°C. Rich or synthetic medium containing 2% lactate, 3% glycerol, 0.05% glucose (YPLGg) and lacking the appropriate amino acids were used to grow the cells overnight prior the induction of I-SceI by plating onto 2% galactose plates or addition of 2% galactose to liquid culture.

### NHEJ efficiency measurement

NHEJ efficiency measurement upon induction of a single DSB was performed as previously described (Batté et al. 2017). Briefly, yeast strains were grown overnight in glycerol lactate containing medium and plated on 2% galactose plates and on 2% glucose plates to respectively induce or repress I-SceI. Survival on galactose was normalized with the cell plating efficiency inferred from survival on glucose. Forty-eight isolated survivors from galactose-containing plates were analysed by PCR and sequenced to characterize NHEJ repair events. For each strain, at least three independent experiments were performed with the corresponding controls.

For plasmid rejoining assays, 50 ng of a pRS316 vector restricted with Xho I was transformed into cells by the lithium acetate transformation method in the presence of 50 µg of denatured salmon sperm DNA as carrier DNA. The number of colonies formed after 3 days was normalized with the number of colonies obtained in a parallel transformation with a circular pRS316 plasmid.

### Monitoring of DSB-flanking DNA and resection by real-time PCR

Yeast cells were grown in 2 mL of YPD overnight. Cultures were then diluted in YPLGg and grown to OD600 = 0.3–0.8.The expression of I-SceI was induced by addition of galactose to a final concentration of 2%. Cell samples were collected before and after induction at different time points and DNAs were extracted. DNA measurements by quantitative PCRs were performed using primers located 0.9 kb from the I-SceI cutting site or primers flanking the I-SceI restriction site. A control primer pair was used to amplify a region of the *OGG1* control locus. To correct for differences in DSB cleavage efficiency, the fraction of uncut DNA (Fu) was subtracted from the fraction of total DNA at 1 kb (Ft) at each time point and normalized to the fraction of cleaved DNA (Fc). Thus, cleaved remaining DNA at 1 kb = (Ft- Fu)/Fc.

### Microscopy

Live cell images were acquired using a wide-field inverted microscope (Leica DMI-6000B) equipped with Adaptive Focus Control to eliminate Z drift, a 100×/1.4 NA immersion objective with a Prior NanoScanZ Nanopositioning Piezo Z Stage System, a CMOS camera (ORCA-Flash4.0; Hamamatsu) and a solid-state light source (SpectraX, Lumencore). The system is piloted by MetaMorph software (Molecular Device).

For GFP-mCherry two-colour images, 19 focal steps of 0.20 μm were acquired sequentially for GFP and mCherry with an exposure time of 100-200 ms using solid-state 475- and 575- nm diodes and appropriate filters (GFP-mCherry filter; excitation: double BP, 450–490/550– 590 nm and dichroic double BP 500–550/600–665 nm; Chroma Technology Corp.). Processing was achieved using ImageJ software (National Institutes of Health). YFP images were acquired at indicated time points before and after DSB induction; 19 focal steps of 0.20 μm were acquired with an exposure time of 200 ms using a solid-state 500-nm diode and a YFP filter (excitation 470–510 nm and dichroic 495 nm; Chroma Technology Corp.) All the images shown are z projections of z-stack images.

### Two-hybrid analyses

The yeast strain Y190 (Wade Harper et al. 1993) was transformed with 2µ plasmids encoding full length or truncated *SAE2* or *SIR3* fused to *GAL4* DNA binding (GBD) or activation (GAD) domains, and selected on synthetic media without leucine and tryptophane. Protein-protein interactions were assayed by growing the cells on selective media without leucine, tryptophane and histidine, complemented with varying concentrations of 3-Amino-1,2,4-triazole (3-AT), a competitive inhibitor of the *HIS3* gene. Blue coloration of the colony in presence of X-Gal was used to assess protein interactions. The interactions were defined in comparison to negative controls, carrying at least one empty vector. When the growth on 3AT containing medium was higher, or if the blue colour in presence of X-gal was stronger than the negative control then an interaction between the two chimeric proteins was assumed.

To screen for *SIR3* mutants a yeast strain (yKD1991) containing *LYS2::GAL1UAS-HIS3TATA-HIS3* and *URA3::lexAop-lacZ* was constructed by crossing Y190 and CTY10-5d (Bartel and Fields 1995). This strain was transformed with 2µ plasmids encoding mutagenized *SIR3^SaID^* fused to *GAL4* DNA binding domain (GBD), *SAE2^C^* fused *LexA* DNA binding domain (LexABD) and *SIR4^C^* to *GAL4* activation domain (GAD).

### Protein fragments cloning and purification

*The Sir3^SaID^* and Sae2^C^ peptides were expressed in *E. coli* strains BL21 (DE3) transformed with pKD434 and pKD435 respectively. Expression of the peptides was induced by 0.5 mM isopropyl-ß-D-thiogalactoside (IPTG) for 3.5 h. Cells were harvested, suspended in lysis buffer (50 mM Tris HCl pH8, 500 mM NaCl, 1 mM DTT, 10% glycerol, TritonX-100 x 1, 1 mg/mL lysozyme, 1 mM 4-(2-aminoethyl) benzenesulphonyl fluoride, 10 mM benzaminide, 2 µM pepstatin) and disrupted by sonication. Extract was cleared by centrifugation at 186000 x g for 1 hour at 4°C.

*Sir3^SaID^* containing extract was incubated at 4°C with GSH Sepharose resin (Cytiva, Marlborough, MA) for 3h. Proteins were eluted with Buffer A (50 mM Tris HCl [pH8@4°C], 100 mM NaCl, 1 mM DTT) complemented with 30 mM glutathione. Fractions containing GST-protein were pooled and applied to a 1 mL Resource Q column (Cytiva, Marlborough, MA) equilibrated with buffer A. Protein was eluted with a 12 mL linear gradient of 0.05–1 M NaCl. Purified GST-protein was stored at -80°C (see purification procedure scheme in Appendix Fig S3).

*Sae2^C^ extract* was incubated with 2mL NiNTA resin (Qiagen, Germantown, MD) in batch, rotated at 4°C for 2h and then poured into a Econo-Column^®^ Chromatography column (Bio-Rad, Hercules, CA). After extensive washing first with 80 mL of 20 mM Tris HCl [pH8@4°C], 500 mM NaCl, 0,5% NP40, 10% glycerol, 20 mM Imidazole followed by 80 mL of 20 mM Tris HCl [pH8@4°C], 100 mM NaCl, 1 mM DTT, 10% glycerol, 20 mM Imidazole, on-column cleavage was achieved by adding his-SUMO-Protease to a ratio of 80/1 (W/W). Untagged *Sae2^Cter^* was recovered from the flow through which was then applied to a 1 mL Resource S column (Cytiva, Marlborough, MA) equilibrated with buffer B (20 mM Tris HCl [pH8@4°C], 50 mM NaCl, 1 mM DTT). Protein was eluted with a 20 mL linear gradient of 0.05–1 M NaCl. Purified *Sae2^Cter^* was stored at -80°C (see purification procedure scheme in Appendix Fig S3).

### GST pull-down assays

GST-*Sir3^SaID^* fragment (10 µg) or GST protein as a control (10µg) were immobilized on 20 µL Glutathione Sepharose 4B in 300 µL of buffer A (50 mM Tris HCl [pH8@4°C], 150 mM NaCl, 1 mM DTT, 0.5 mM EDTA, 10% Glycerol), complemented with 2 mM MgCl_2_ and 25 units of benzonase for 90 minutes at 4°C. Beads were collected by centrifugation, and washed three times with 300 µL of buffer B (buffer A + 0.05% NP40). *Sae2^C^* (10 µg in 100 µL buffer B complemented with 2 mM MgCl_2_ and 25 units of benzonase) was then added and incubated for 150 minutes at 4°C with gentle agitation. The supernatant was removed and the beads were washed two times with 300 µL of buffer B. Proteins bound to the beads were then eluted by addition of 20 µL of 50 mM Tris-HCl [pH8@4°C], 150 mM NaCl, 1 mM DTT, 30 mM glutathione. Proteins bound to the beads were resolved on 15% SDS-PAGE.

### Co-immunoprecipitation (Co-IP)

Immunoprecipitations were performed as previously described (Forey et al. 2021) by lysing 40 to 50 OD600 units of exponential phase cultures. After sonication, clarification and benzonase treatment (250 u/1 mg protein, SIGMA E1014-5KU) extracts were incubated 1h at 4°C with 50 µL magnetic beads (Dynabeads M-280 sheep anti-mouse IgG ,invitrogen 11202D) coated with anti-GFP antibodies (Roche ref:11814460001). Proteins extracts were resolved on 4-15% polyacrylamide gels, transferred on iBlot PVDF Membranes that were probed with anti-GFP (1:1000, Roche ref:11814460001) and custom-made anti-Sir3 (1:10000;(Ruault et al. 2011)) antibodies.

### Chromatin immunoprecipitation (ChIP)

Exponentially growing cells were crosslinked for 15 min with 1% formaldehyde (Sigma F8775) at RT under agitation followed by quenching by addition of 0.125 M Glycine (Sigma G8898) for 5 min under agitation. Cells were washed three times with cold 20mM Tris (4°C). Dry pellets were frozen and conserved at -80°C. Cell pellets were resuspended in lysis buffer (50 mM HEPES-KOH pH7.5, 140 mM NaCl, 5 mM EDTA, 1% Triton X-100, 0.1% Na-deoxycholate) supplemented with 1 mM AEBSF (ThermoFisher 10563165) and anti-protease (Complete ULTRA SIGMA ref: 5892988001) and lysed with a Precellys homogenizer. Whole cell extracts were centrifuged 20 min at 13535 rpm and the chromatin containing pellet was resuspended in 300 µL lysis buffer. Sonication of chromatin was performed using a Diagenod Bioruptor at high setting for 3 cycles: 30 seconds ON + 30 seconds OFF. Dynabeads (Panmous IgG, Invitrogen 11041) were washed three times and resuspended in 1 mL of PBS, 0.1% BSA and incubated with antibodies (10 µL anti-GFP (1:1000, Roche ref:11814460001)/50 µL beads) on a rotating wheel for two hours at 4 °C. Antibody-coupled Dynabeads were washed three times with 1 mL of PBS, 0.1% BSA, and incubated with 400 µL of sonicated chromatin for 2h at 21°C. Beads were washed on ice with cold solutions: two times with lysis buffer, once with wash buffer (10 mM Tris-HCl pH8, 0.25 M LiCl, 0.5% NP40, 5 mM EDTA, 0.5% Na-deoxycholate) and once with TE (10 mM Tris-HCl pH8, 1 mM EDTA). Antibodies were un-coupled from beads with elution buffer (25 mM Tris-HCl pH8, 5 mM EDTA, 0,5% SDS) for 20 min at 65 °C. Eluates were collected and incubated overnight at 65°C for de-crosslinking. RNAse A (Sigma, R65-13) and Pronase were added to samples and incubated for 1 hour at 37 °C. DNA was purified (DNA clean up kit, Thermoscientific K0832) and eluted in 50 µl of elution buffer. The relative amount of DNA was quantified by qPCR (primers listed in Appendix Table S2). Sae2-GFP enrichment was normalized to an internal control locus (OGG1).

### Data availability

This study includes no data deposited in external repositories.

## Acknowledgements

We thank Z. Xu, A. Piazza, R. Koszul and Jamie Phipps for critical reading of this manuscript and members of the Marcand and Dubrana laboratories for stimulating discussions. We thank Susan Gasser, Ann E. Ehrenhofer-Murray, Lorraine Symington, Rodney Rothstein, Laurent Maloisel and Florian Roisné-Hamelin for sharing plasmids and strains and Angela Taddei for the custom-made rabbit polyclonal antibodies raised against full-length Sir3. This work was supported by grants from Fondation pour la recherche médicale (DEP20131128535), and from the European Research Council under the European Community’s Seventh Frame-work Program (FP7/2007 2013/European Research Council Grant Agreement 281287), Fondation ARC pour la Recherche sur le Cancer (PJA-20191209432), CEA Radiation biology program and EDF. HB was supported by a fellowship from the CEA-IRTELIS PhD program.

## Author contributions

HB, RC, CB, AB and KD performed experiments in yeast. DB, JD and XV performed the *in vitro* experiments. SM designed and supervised the telomere length experiments and gave critical input on NHEJ experiments. RG performed protein alignments that helped design protein subdomains for yeast two hybrid and expression in bacteria. KD designed and supervised the entire project with the help of HB. KD and HB wrote the manuscript with critical input of the other authors.

## Expanded View Figure Legends

**Figure EV1:**
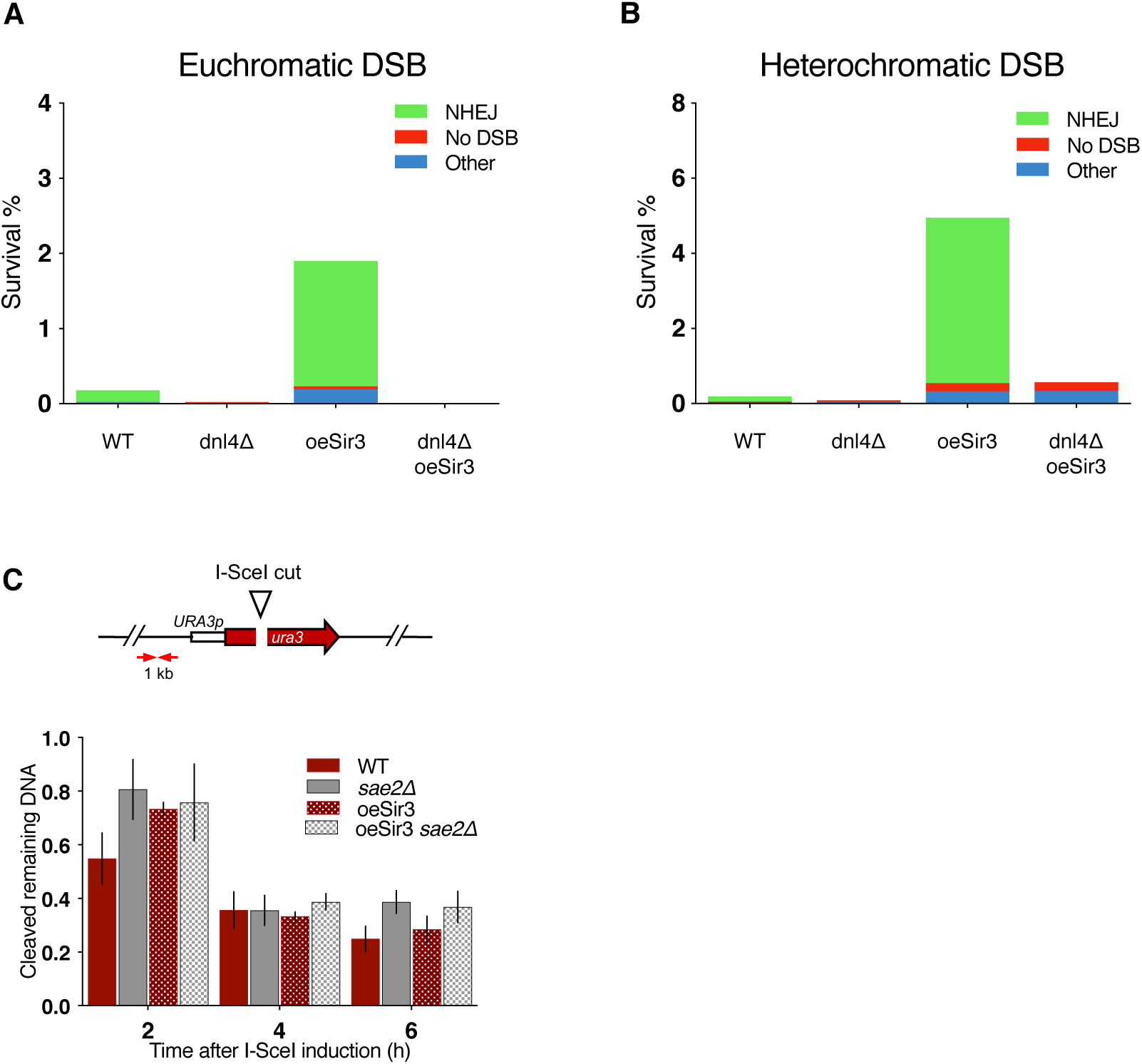
A. Survival frequencies and characterisation of the repair events after induction of a DSB at LYS2 in absence of recombination substrate. B. Survival frequencies and characterisation of the repair events after induction of a DSB at TEL6R in absence of recombination substrate. NHEJ stands for error-prone end joining events detected by a PCR product that cannot be cleaved *in vitro* by I-SceI. No DSB corresponds to survivors giving a PCR product that can be cleaved by I-SceI *in vitro* showing that they failed to induce I-SceI . Other gathers survivors in which no PCR product was obtained suggesting that repair occured through other mechanisms. PCR products corresponding to NHEJ events were sequenced and exhibit patterns typical of NHEJ repair (rejoining with 1 to 9 bp deletion between sequences showing no or limited homology). C. DNA levels measured at 1 kb from the I-SceI cut site at LYS2 after 2h DSB induction by qPCR in WT and sae2Δ cells expressing or not high levels of Sir3p (oeSir3 and WT respectively). DNA levels were normalized to DNA levels at the OGG1 locus and corrected for differences in DSB cleavage efficiency (see Materials and Methods for details). Error bars represent the standard deviation (SD) of three independent experiments.

**Figure EV2:**
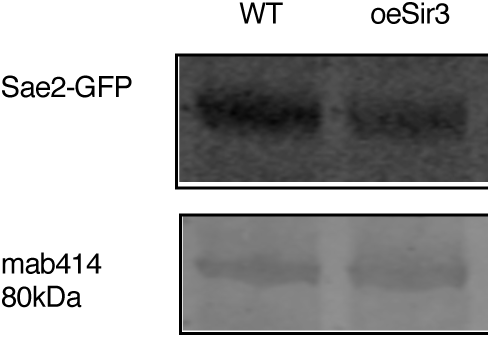
Western blot analysis with anti-GFP antibodies of whole cell protein extracts prepared from stationary phase cells. The 80kDa band detected by mab414 is used as a loading control.

**Figure EV3:**
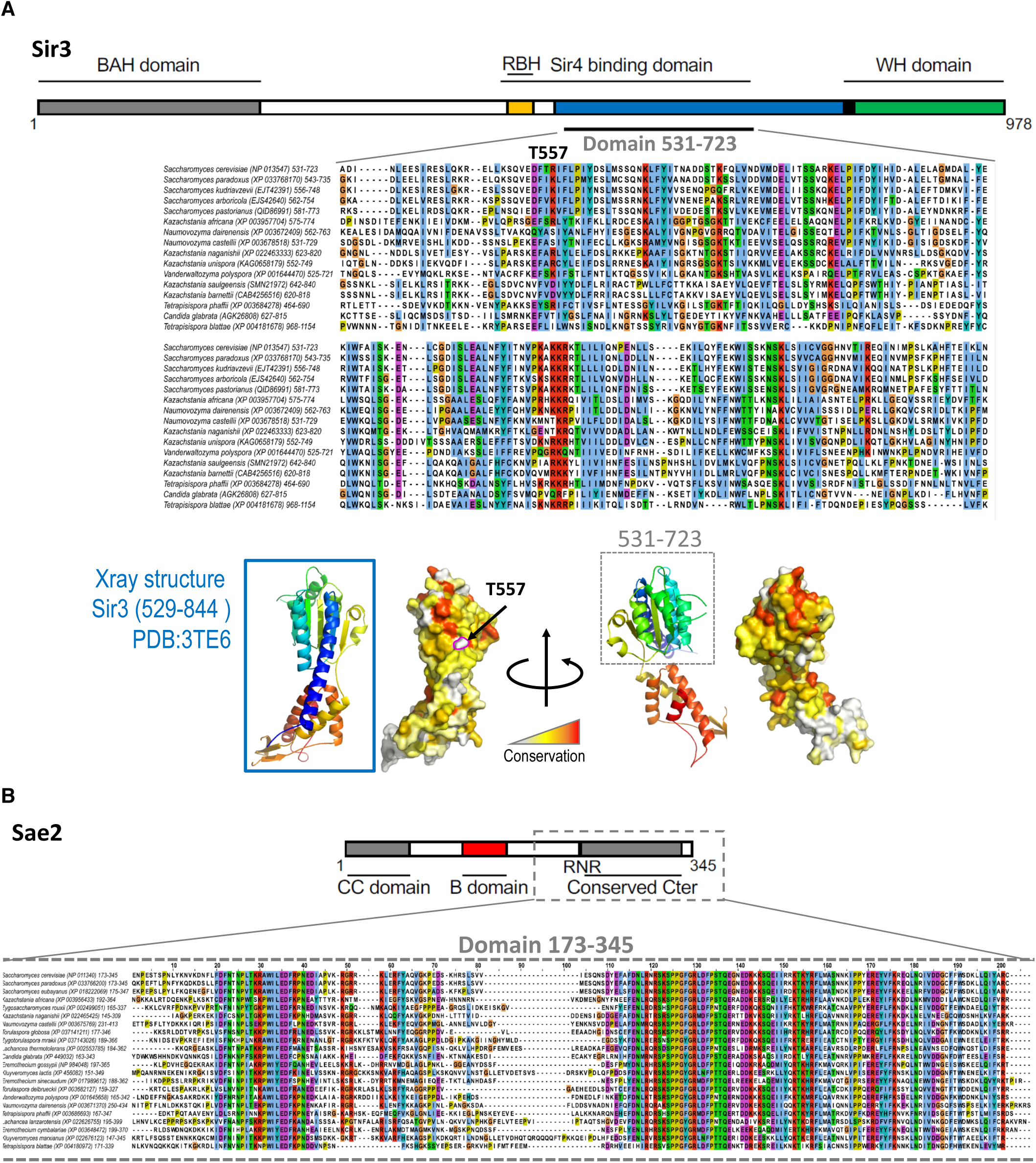
A. Multiple sequence alignment of the Sir3^SaID^domain of the *Saccharomycetaceae* family. NCBI RefSeq identifiers are given in parentheses. Below, ribon representation of the Xray structure of the Sir3^SaID^domain (rainbow colors) and a surface projection of the conservation as calculated by the rate4site algorithm (Pupko et al, 2002) with a white-yellow-red color gradient highlighting the most conserved region in red. B. Multiple sequence alignment of the Sae2^C^domain of the *Saccharomycetaceae* family. NCBI RefSeq identifiers are given in parentheses.

**Figure EV4:**
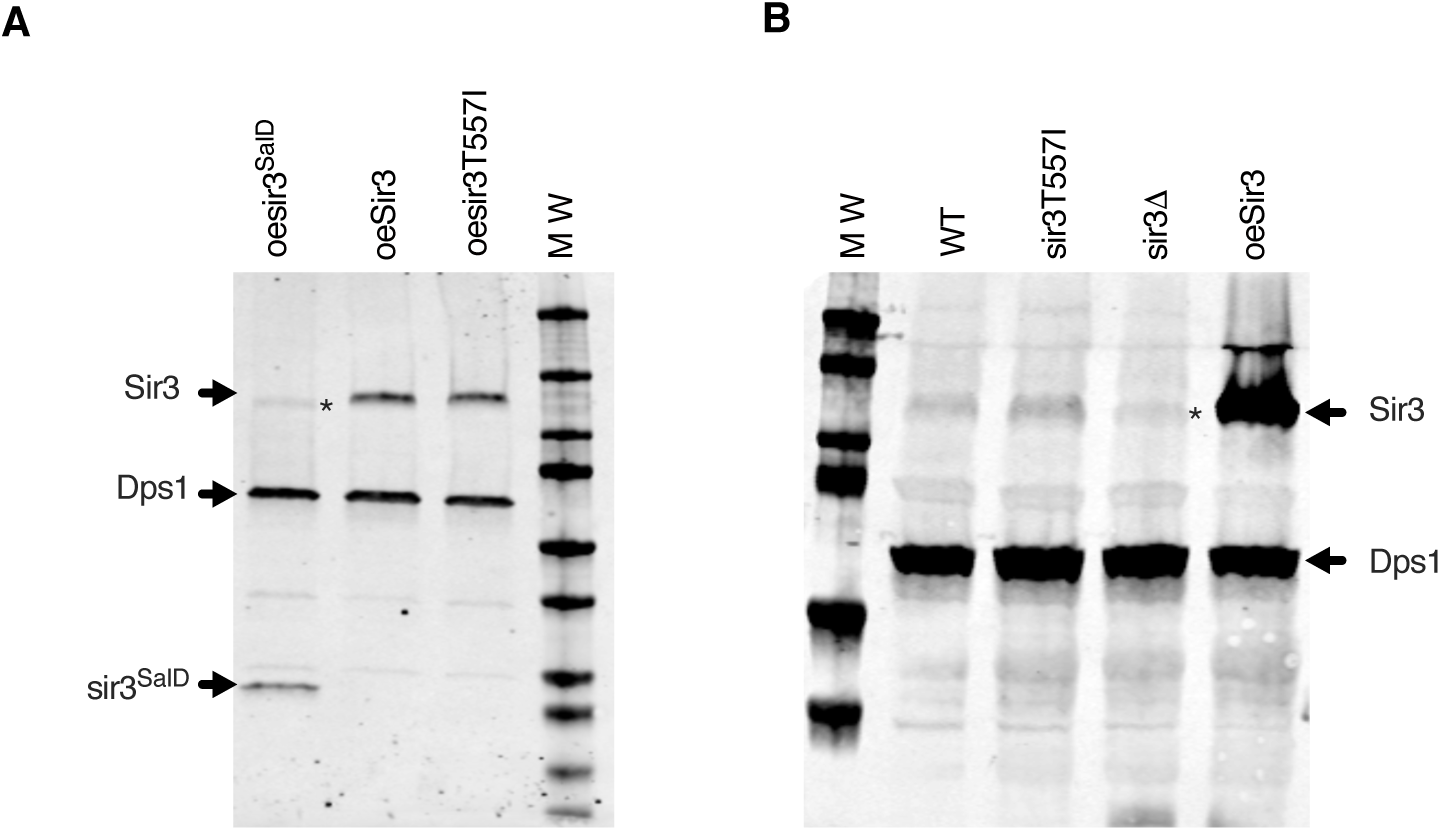
A, B. Western blot analysis with Sir3 antibodies of protein extracts prepared from stationary phase cells of the indicated strains. Dps1 is used as a loading control. *Asterisk marks cross reacting Orc1 detected by the Sir3 antibody.

**Figure EV5:**
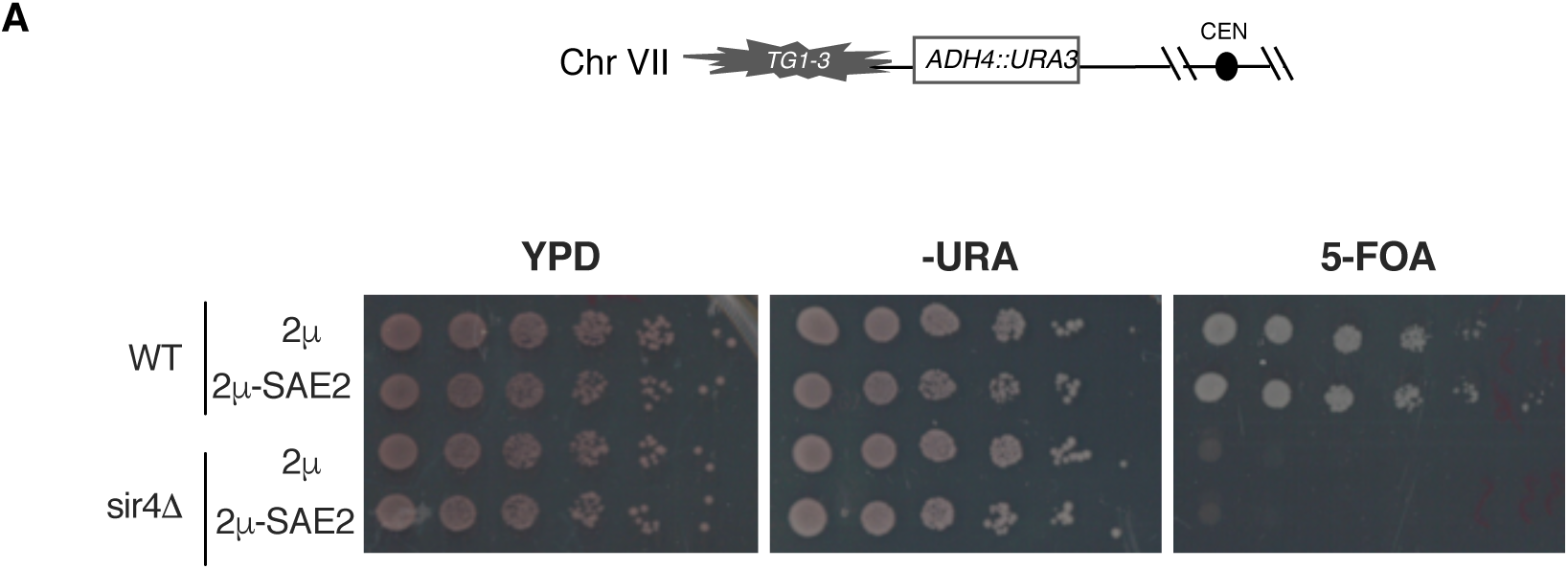
A. Telomeric silencing assay at TEL7L in WT and sir4Δ cells overexpressing SAE2 (2μ-Sae2) or not (2 μ). Increased growth on 5-FOA or decreased growth on -URA plates reflects an increase in telomeric silencing.

## References

Allshire RC, Madhani HD (2018) Ten principles of heterochromatin formation and function. Nat Rev Mol Cell Biol 19:229–244. https://doi.org/10.1038/nrm.2017.119

Arnerić M, Lingner J (2007) Tel1 kinase and subtelomere-bound Tbf1 mediate preferential elongation of short telomeres by telomerase in yeast. EMBO Rep 8:1080–1085. https://doi.org/10.1038/sj.embor.7401082

Aström SU, Okamura SM, Rine J (1999) Yeast cell-type regulation of DNA repair. Nature 397:310. https://doi.org/10.1038/16833

Aymard F, Bugler B, Schmidt CK, et al (2014) Transcriptionally active chromatin recruits homologous recombination at DNA double-strand breaks. Nat Struct Mol Biol 21:366–374. https://doi.org/10.1038/nsmb.2796

Baroni E, Viscardi V, Cartagena-Lirola H, et al (2004) The Functions of Budding Yeast Sae2 in the DNA Damage Response Require Mec1-and Tel1-Dependent Phosphorylation. Mol Cell Biol 24:4151–4165. https://doi.org/10.1128/MCB.24.10.4151-4165.2004

Bartel PL, Fields S (1995) Analyzing protein-protein interactions using two-hybrid system. Methods Enzymol 254:241–263. https://doi.org/10.1016/0076-6879(95)54018-0

Batté A, Brocas C, Bordelet H, et al (2017) Recombination at subtelomeres is regulated by physical distance, double-strand break resection and chromatin status. EMBO J 36:2609–2625. https://doi.org/10.15252/embj.201796631

Bazzano D, Lomonaco S, Wilson TE (2021) Mapping yeast mitotic 5’ resection at base resolution reveals the sequence and positional dependence of nucleases in vivo. bioRxiv 2021.02.27.433206. https://doi.org/10.1101/2021.02.27.433206

Behrouzi R, Lu C, Currie MA, et al (2016) Heterochromatin assembly by interrupted Sir3 bridges across neighboring nucleosomes. eLife 5:. https://doi.org/10.7554/eLife.17556

Bonetti D, Martina M, Clerici M, et al (2009) Multiple pathways regulate 3’ overhang generation at S. cerevisiae telomeres. Mol Cell 35:70–81. https://doi.org/10.1016/j.molcel.2009.05.015

Cannavo E, Cejka P (2014) Sae2 promotes dsDNA endonuclease activity within Mre11-Rad50-Xrs2 to resect DNA breaks. Nature 514:122–125. https://doi.org/10.1038/nature13771

Cannavo E, Johnson D, Andres SN, et al (2018) Regulatory control of DNA end resection by Sae2 phosphorylation. Nat Commun 9:4016. https://doi.org/10.1038/s41467-018-06417-5

Chen H, Lisby M, Symington LS (2013) RPA coordinates DNA end resection and prevents formation of DNA hairpins. Mol Cell 50:. https://doi.org/10.1016/j.molcel.2013.04.032

Chen H, Xue J, Churikov D, et al (2018) Structural Insights into Yeast Telomerase Recruitment to Telomeres. Cell 172:331–343.e13. https://doi.org/10.1016/j.cell.2017.12.008

Chen X, Tomkinson AE (2011) Yeast Nej1 Is a Key Participant in the Initial End Binding and Final Ligation Steps of Nonhomologous End Joining. J Biol Chem 286:4931–4940. https://doi.org/10.1074/jbc.M110.195024

Chiolo I, Minoda A, Colmenares SU, et al (2011) Double-strand breaks in heterochromatin move outside of a dynamic HP1a domain to complete recombinational repair. Cell 144:732–744. https://doi.org/10.1016/j.cell.2011.02.012

Clerici M, Mantiero D, Lucchini G, Longhese MP (2006) The Saccharomyces cerevisiae Sae2 protein negatively regulates DNA damage checkpoint signalling. EMBO Rep 7:212–218. https://doi.org/10.1038/sj.embor.7400593

Cubizolles F, Martino F, Perrod S, Gasser SM (2006) A homotrimer-heterotrimer switch in Sir2 structure differentiates rDNA and telomeric silencing. Mol Cell 21:825– 836. https://doi.org/10.1016/j.molcel.2006.02.006

Dalby AB, Goodrich KJ, Pfingsten JS, Cech TR (2013) RNA recognition by the DNA end-binding Ku heterodimer. RNA 19:841–851. https://doi.org/10.1261/rna.038703.113

Diebold M-L, Fribourg S, Koch M, et al (2011) Deciphering correct strategies for multiprotein complex assembly by co-expression: application to complexes as large as the histone octamer. J Struct Biol 175:178–188. https://doi.org/10.1016/j.jsb.2011.02.001

Ehrentraut S, Hassler M, Oppikofer M, et al (2011) Structural basis for the role of the Sir3 AAA+ domain in silencing: interaction with Sir4 and unmethylated histone H3K79. Genes Dev 25:1835–1846. https://doi.org/10.1101/gad.17175111

Faure G, Jézéquel K, Roisné-Hamelin F, et al (2019) Discovery and Evolution of New Domains in Yeast Heterochromatin Factor Sir4 and Its Partner Esc1. Genome Biol Evol 11:572–585. https://doi.org/10.1093/gbe/evz010

Forey R, Barthe A, Tittel-Elmer M, et al (2021) A Role for the Mre11-Rad50-Xrs2 Complex in Gene Expression and Chromosome Organization. Mol Cell 81:183–197.e6. https://doi.org/10.1016/j.molcel.2020.11.010

Frank-Vaillant M, Marcand S (2001) NHEJ regulation by mating type is exercised through a novel protein, Lif2p, essential to the Ligase IV pathway. Genes Dev 15:3005–3012. https://doi.org/10.1101/gad.206801

Frank-Vaillant M, Marcand S (2002) Transient Stability of DNA Ends Allows Nonhomologous End Joining to Precede Homologous Recombination. Mol Cell 10:1189–1199. https://doi.org/10.1016/S1097-2765(02)00705-0

Fu Q, Chow J, Bernstein KA, et al (2014) Phosphorylation-Regulated Transitions in an Oligomeric State Control the Activity of the Sae2 DNA Repair Enzyme. Mol Cell Biol 34:778–793. https://doi.org/10.1128/MCB.00963-13

Garcia V, Phelps SEL, Gray S, Neale MJ (2011) Bidirectional resection of DNA double-strand breaks by Mre11 and Exo1. Nature 479:241. https://doi.org/10.1038/nature10515

Gartenberg MR, Smith JS (2016) The Nuts and Bolts of Transcriptionally Silent Chromatin in Saccharomyces cerevisiae. Genetics 203:1563–1599. https://doi.org/10.1534/genetics.112.145243

Goodarzi AA, Noon AT, Deckbar D, et al (2008) ATM Signaling Facilitates Repair of DNA Double-Strand Breaks Associated with Heterochromatin. Mol Cell 31:167–177. https://doi.org/10.1016/j.molcel.2008.05.017

Hansson MD, Rzeznicka K, Rosenbäck M, et al (2008) PCR-mediated deletion of plasmid DNA. Anal Biochem 375:373–375. https://doi.org/10.1016/j.ab.2007.12.005

Hass EP, Zappulla DC (2015) The Ku subunit of telomerase binds Sir4 to recruit telomerase to lengthen telomeres in S. cerevisiae. In: eLife. https://elifesciences.org/articles/07750. Accessed 20 Apr 2021

Hecht A, Strahl-Bolsinger S, Grunstein M (1996) Spreading of transcriptional represser SIR3 from telomeric heterochromatin. Nature 383:92. https://doi.org/10.1038/383092a0

Hocher A, Ruault M, Kaferle P, et al (2018) Expanding heterochromatin reveals discrete subtelomeric domains delimited by chromatin landscape transitions. Genome Res 28:1867–1881. https://doi.org/10.1101/gr.236554.118

Huertas P, Cortés-Ledesma F, Sartori AA, et al (2008) CDK targets Sae2 to control DNA-end resection and homologous recombination. Nature 455:689–692. https://doi.org/10.1038/nature07215

Huertas P, Jackson SP (2009) Human CtIP mediates cell cycle control of DNA end resection and double strand break repair. J Biol Chem 284:9558–9565. https://doi.org/10.1074/jbc.M808906200

Kaiser GS, Germann SM, Westergaard T, Lisby M (2011) Phenylbutyrate inhibits homologous recombination induced by camptothecin and methyl methanesulfonate. Mutat Res 713:64–75. https://doi.org/10.1016/j.mrfmmm.2011.05.016

Katan-Khaykovich Y, Struhl K (2005) Heterochromatin formation involves changes in histone modifications over multiple cell generations. EMBO J 24:2138–2149. https://doi.org/10.1038/sj.emboj.7600692

Kegel A, Sjöstrand JOO, Åström SU (2001) Nej1p, a cell type-specific regulator of nonhomologous end joining in yeast. Curr Biol 11:1611–1617. https://doi.org/10.1016/S0960-9822(01)00488-2

Khayat F, Cannavo E, Alshmery M, et al (2021) Inhibition of MRN activity by a telomere protein motif. Nat Commun 12:3856. https://doi.org/10.1038/s41467-021-24047-2

King DA, Hall BE, Iwamoto MA, et al (2006) Domain Structure and Protein Interactions of the Silent Information Regulator Sir3 Revealed by Screening a Nested Deletion Library of Protein Fragments. J Biol Chem 281:20107–20119. https://doi.org/10.1074/jbc.M512588200

Larson AG, Elnatan D, Keenen MM, et al (2017) Liquid droplet formation by HP1α suggests a role for phase separation in heterochromatin. Nature 547:236–240. https://doi.org/10.1038/nature22822

Lee C-S, Wang RW, Chang H-H, et al (2016) Chromosome position determines the success of double-strand break repair. Proc Natl Acad Sci U S A 113:E146–154. https://doi.org/10.1073/pnas.1523660113

Lee K, Lee SE (2007) Saccharomyces cerevisiae Sae2- and Tel1-Dependent Single-Strand DNA Formation at DNA Break Promotes Microhomology-Mediated End Joining. Genetics 176:2003–2014. https://doi.org/10.1534/genetics.107.076539

Lee SE, Pâques F, Sylvan J, Haber JE (1999) Role of yeast SIR genes and mating type in directing DNA double-strand breaks to homologous and non-homologous repair paths. Curr Biol CB 9:767–770

Lemaître C, Grabarz A, Tsouroula K, et al (2014) Nuclear position dictates DNA repair pathway choice. Genes Dev 28:2450–2463. https://doi.org/10.1101/gad.248369.114

Léon S, Erpapazoglou Z, Haguenauer-Tsapis R (2008) Ear1p and Ssh4p Are New Adaptors of the Ubiquitin Ligase Rsp5p for Cargo Ubiquitylation and Sorting at Multivesicular Bodies. Mol Biol Cell 19:2379–2388. https://doi.org/10.1091/mbc.E08-01-0068

Li MZ, Elledge SJ (2007) Harnessing homologous recombination in vitro to generate recombinant DNA via SLIC. Nat Methods 4:251–256. https://doi.org/10.1038/nmeth1010

Liang J, Suhandynata RT, Zhou H (2015) Phosphorylation of Sae2 Mediates Forkhead-associated (FHA) Domain-specific Interaction and Regulates Its DNA Repair Function. J Biol Chem 290:10751–10763. https://doi.org/10.1074/jbc.M114.625293

Lisby M, Rothstein R (2004) Choreography of the DNA damage response: spatiotemporal relationships among checkpoint and repair proteins. Cell 118:699– 713. https://doi.org/10.1016/j.cell.2004.08.015

Longtine MS, McKenzie A, Demarini DJ, et al (1998) Additional modules for versatile and economical PCR-based gene deletion and modification in Saccharomyces cerevisiae. Yeast Chichester Engl 14:953–961. https://doi.org/10.1002/(SICI)1097-0061(199807)14:10<953::AID-YEA293>3.0.CO;2-U

Machida S, Takizawa Y, Ishimaru M, et al (2018) Structural Basis of Heterochromatin Formation by Human HP1. Mol Cell 69:385–397.e8. https://doi.org/10.1016/j.molcel.2017.12.011

Mahaney BL, Lees-Miller SP, Cobb JA (2014) The C-terminus of Nej1 is critical for nuclear localization and non-homologous end-joining. DNA Repair 14:9–16. https://doi.org/10.1016/j.dnarep.2013.12.002

Marcand S, Pardo B, Gratias A, et al (2008) Multiple pathways inhibit NHEJ at telomeres. Genes Dev 22:1153–1158. https://doi.org/10.1101/gad.455108

Martin SG, Laroche T, Suka N, et al (1999) Relocalization of Telomeric Ku and SIR Proteins in Response to DNA Strand Breaks in Yeast. Cell 97:621–633. https://doi.org/10.1016/S0092-8674(00)80773-4

Matsuzaki K, Shinohara A, Shinohara M (2008) Forkhead-Associated Domain of Yeast Xrs2, a Homolog of Human Nbs1, Promotes Nonhomologous End Joining Through Interaction With a Ligase IV Partner Protein, Lif1. Genetics 179:213–225. https://doi.org/10.1534/genetics.107.079236

McAinsh AD, Scott-Drew S, Murray JA, Jackson SP (1999) DNA damage triggers disruption of telomeric silencing and Mec1p-dependent relocation of Sir3p. Curr Biol CB 9:963–966

Meister P, Taddei A (2013) Building silent compartments at the nuclear periphery: a recurrent theme. Curr Opin Genet Dev 23:96–103. https://doi.org/10.1016/j.gde.2012.12.001

Mills KD, Sinclair DA, Guarente L (1999) MEC1-Dependent Redistribution of the Sir3 Silencing Protein from Telomeres to DNA Double-Strand Breaks. Cell 97:609–620. https://doi.org/10.1016/S0092-8674(00)80772-2

Mimitou EP, Symington LS (2008) Sae2, Exo1 and Sgs1 collaborate in DNA double-strand break processing. Nature 455:770–774. https://doi.org/10.1038/nature07312

Mojumdar A, Adam N, Cobb JA (2021) Sgs1BLM independent role of Dna2DNA2 nuclease at DNA double strand break is inhibited by Nej1XLF

Muraki K, Han L, Miller D, Murnane JP (2015) Processing by MRE11 is involved in the sensitivity of subtelomeric regions to DNA double-strand breaks. Nucleic Acids Res 43:7911–7930. https://doi.org/10.1093/nar/gkv714

Palmbos PL, Daley JM, Wilson TE (2005) Mutations of the Yku80 C Terminus and Xrs2 FHA Domain Specifically Block Yeast Nonhomologous End Joining. Mol Cell Biol 25:10782–10790. https://doi.org/10.1128/MCB.25.24.10782-10790.2005

Palmbos PL, Wu D, Daley JM, Wilson TE (2008) Recruitment of Saccharomyces cerevisiae Dnl4–Lif1 Complex to a Double-Strand Break Requires Interactions With Yku80 and the Xrs2 FHA Domain. Genetics 180:1809–1819. https://doi.org/10.1534/genetics.108.095539

Renauld H, Aparicio OM, Zierath PD, et al (1993) Silent domains are assembled continuously from the telomere and are defined by promoter distance and strength, and by SIR3 dosage. Genes Dev 7:1133–1145. https://doi.org/10.1101/gad.7.7a.1133

Robert T, Vanoli F, Chiolo I, et al (2011) HDACs link the DNA damage response, processing of double-strand breaks and autophagy. Nature 471:74–79. https://doi.org/10.1038/nature09803

Roisné-Hamelin F, Pobiega S, Jézéquel K, et al (2021) Mechanism of MRX inhibition by Rif2 at telomeres. Nat Commun 12:2763. https://doi.org/10.1038/s41467-021-23035-w

Roy R, Meier B, McAinsh AD, et al (2004) Separation-of-function Mutants of Yeast Ku80 Reveal a Yku80p-Sir4p Interaction Involved in Telomeric Silencing. J Biol Chem 279:86–94. https://doi.org/10.1074/jbc.M306841200

Ruault M, De Meyer A, Loïodice I, Taddei A (2011) Clustering heterochromatin: Sir3 promotes telomere clustering independently of silencing in yeast. J Cell Biol 192:417–431. https://doi.org/10.1083/jcb.201008007

Ruault M, Scolari VF, Lazar-Stefanita L, et al (2021) Sir3 mediates long-range chromosome interactions in budding yeast. Genome Res 31:411–425. https://doi.org/10.1101/gr.267872.120

Sheff MA, Thorn KS (2004) Optimized cassettes for fluorescent protein tagging in Saccharomyces cerevisiae. Yeast Chichester Engl 21:661–670. https://doi.org/10.1002/yea.1130

Soria-Bretones I, Cepeda-García C, Checa-Rodriguez C, et al (2017) DNA end resection requires constitutive sumoylation of CtIP by CBX4. Nat Commun 8:113. https://doi.org/10.1038/s41467-017-00183-6

Strahl-Bolsinger S, Hecht A, Luo K, Grunstein M (1997) SIR2 and SIR4 interactions differ in core and extended telomeric heterochromatin in yeast. Genes Dev 11:83–93. https://doi.org/10.1101/gad.11.1.83

Strom AR, Emelyanov AV, Mir M, et al (2017) Phase separation drives heterochromatin domain formation. Nature 547:241–245. https://doi.org/10.1038/nature22989

Swygert SG, Senapati S, Bolukbasi MF, et al (2018) SIR proteins create compact heterochromatin fibers. Proc Natl Acad Sci 115:12447–12452. https://doi.org/10.1073/pnas.1810647115

Symington LS (2016) Mechanism and regulation of DNA end resection in eukaryotes. Crit Rev Biochem Mol Biol 51:195–212. https://doi.org/10.3109/10409238.2016.1172552

Toledo LI, Altmeyer M, Rask M-B, et al (2013) ATR Prohibits Replication Catastrophe by Preventing Global Exhaustion of RPA. Cell 155:1088–1103. https://doi.org/10.1016/j.cell.2013.10.043

Tsabar M, Eapen VV, Mason JM, et al (2015) Caffeine impairs resection during DNA break repair by reducing the levels of nucleases Sae2 and Dna2. Nucleic Acids Res 43:6889–6901. https://doi.org/10.1093/nar/gkv520

Tsouroula K, Furst A, Rogier M, et al (2016) Temporal and Spatial Uncoupling of DNA Double Strand Break Repair Pathways within Mammalian Heterochromatin. Mol Cell 63:293–305. https://doi.org/10.1016/j.molcel.2016.06.002

Valencia M, Bentele M, Vaze MB, et al (2001) NEJ1 controls non-homologous end joining in Saccharomyces cerevisiae. Nature 414:666–669. https://doi.org/10.1038/414666a

van Steensel B, Smogorzewska A, de Lange T (1998) TRF2 protects human telomeres from end-to-end fusions. Cell 92:401–413

Wade Harper J, Adami GR, Wei N, et al (1993) The p21 Cdk-interacting protein Cip1 is a potent inhibitor of G1 cyclin-dependent kinases. Cell 75:805–816. https://doi.org/10.1016/0092-8674(93)90499-G

Wu W, Nishikawa H, Fukuda T, et al (2015) Interaction of BARD1 and HP1 is required for BRCA1 retention at sites of DNA damage. Cancer Res 75:1311–1321. https://doi.org/10.1158/0008-5472.CAN-14-2796

Yu H, Braun P, Yildirim MA, et al (2008) High-quality binary protein interaction map of the yeast interactome network. Science 322:104–110. https://doi.org/10.1126/science.1158684

Yu T-Y, Kimble MT, Symington LS (2018) Sae2 antagonizes Rad9 accumulation at DNA double-strand breaks to attenuate checkpoint signaling and facilitate end resection. Proc Natl Acad Sci U S A 115:E11961–E11969. https://doi.org/10.1073/pnas.1816539115

Zdravković A, Daley JM, Dutta A, et al (2021) A conserved Ctp1/CtIP C-terminal peptide stimulates Mre11 endonuclease activity. Proc Natl Acad Sci U S A 118:.https://doi.org/10.1073/pnas.2016287118

