## Supplementary material for "Sir3 Heterochromatin Protein Promotes NHEJ by Direct Inhibition of Sae2": EMBOJ-2021-108813_Appendix.pdf

**Table of Contents**

Appendix Figure S1

Appendix Figure S2

Appendix Figure S3

Appendix Table S1

Appendix Table S2

Bordelet et al, Appendix Figure S1

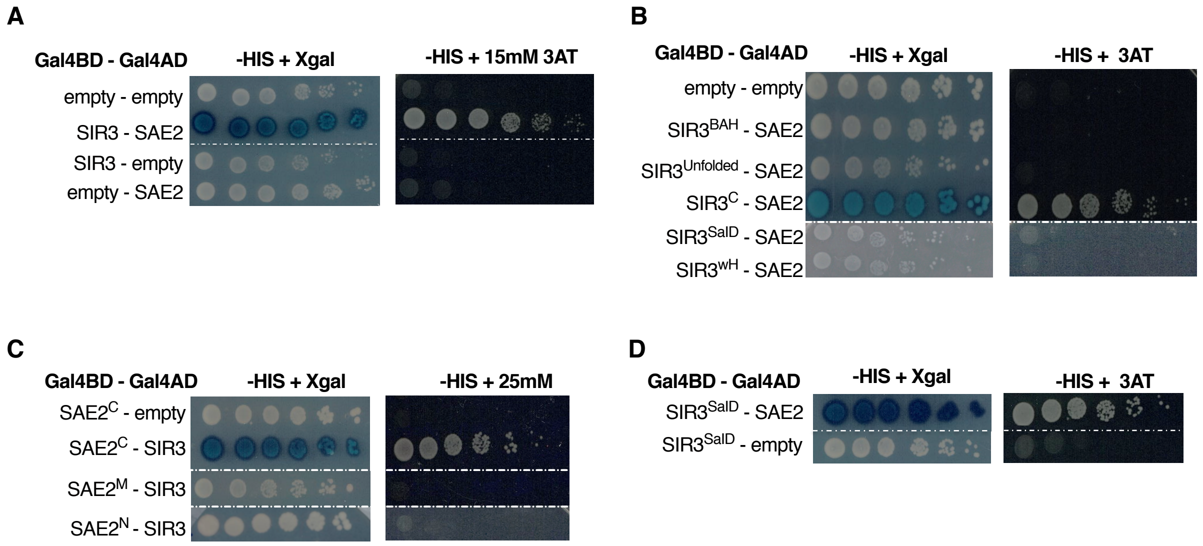


**Yeast two-hybrid analysis of Sir3-Sae2 interaction**

Y2H screen analysis was performed in WT cells on -His + 3AT media and X gal coloration.

A. Representative image of two hybrid assays testing the interaction of the full length WT Sae2 and Sir3 proteins.

B. Representative image of two hybrid assays testing the interaction of full length Sae2 with Sir3 fragments. $SIR3^{BAH}$= aa 1-216, $SIR3^{unfolded}$= aa 217-530 SIR3C= aa 531-848, SIR3wH= aa 849-978 and $SIR3^{SalD}$= 531-723.

C. Representative image of two hybrid assays testing the interaction of full length Sir3 with Sae2 fragments. $SAE2^{C}$= aa 1-72 , $SAE2^{M}$= aa 73-172 , $SAE2^{N}$= aa 173-345.

D. Representative image of two hybrid assays testing the interaction of $Sir3^{SalD}$ and $Sae2^{C}$ with full length Sae2 and Sir3 respectively.

Bordelet et al, Appendix Figure S2

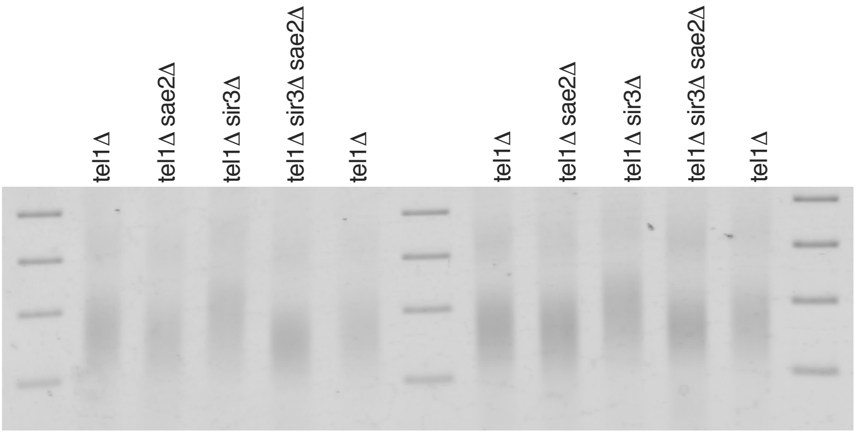


Representative electrophoresis of telomere-PCR products of tel1Δ strains combined with the indicated null mutations. Telomere length was assessed by PCR after end-labaling with terminal transferase (Teixeira et al 2004).

Bordelet et al, Appendix Figure S3

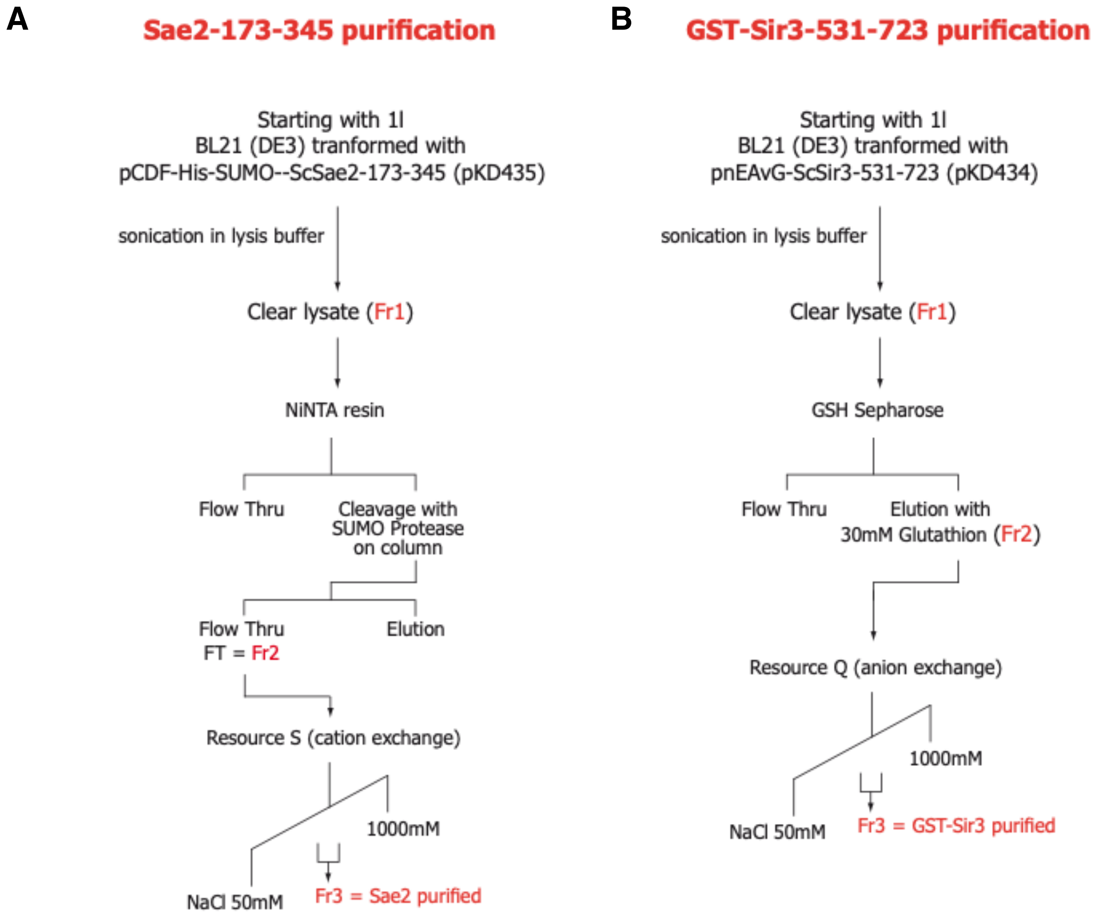


Scheme of the purification procedure for Sae2 (A) and Sir3 (B) fragments

Appendix Table S1**: S. cerevisiae strains used in this paper:**

| **Strain** | | **Genotype** | **Source or reference** | **figure** |
|---|---|---|---|---|
| yKD513 | (1) | TELVI-R::lox-ura3-ISceI ura3-1Δ::KanMx | This study | 1B |
| yKD788 | (1) | TELVI-R::lox-ura3-ISceI ura3-1Δ::KanMx dnl4Δ::HIS3Mx | Batté et al. 2107 | 1B |
| yKD790 | (1) | TELVI-R::lox-ura3-ISceI ura3-1Δ::KanMx sir3::Nat-GPDp-SIR3 | This study | 1B, 2C |
| yKD800 | (1) | TELVI-R::lox-ura3-ISceI ura3-1Δ::KanMx dnl4Δ::HIS3Mx sir3::Nat-GPDp-SIR3 | Batté et al. 2107 | 1B |
| yKD1891 | (1) | TELVI-R::lox-ura3-ISceI ura3-1Δ::KanMx sir3::Nat-GPDp-SIR3 sir4Δ::HIS3Mx | This study | 1B |
| yKD1472 | (1) | TELVI-R::lox-ura3-ISceI ura3-1Δ::KanMx sae2Δ::HIS3Mx | This study | 1B |
| yKD1561 | (1) | TELVI-R::lox-ura3-ISceI ura3-1Δ::KanMx sae2Δ::HIS3Mx sir3::Nat-GPDp-SIR3 | This study | 1B |
| yKD516 | (1) | lys2::lox-ura3-ISceI ura3-1Δ::KanMx | This study | 1C, 1E, 1H, 5A, 5B, 6A, 6B |
| yKD789 | (1) | lys2::lox-ura3-ISceI ura3-1Δ::KanMx dnl4Δ::HIS3Mx | Batté et al. 2107 | 1C, 6B |
| yKD706 | (1) | lys2::lox-ura3-ISceI ura3-1Δ::KanMx sir3::Nat-GPDp-SIR3 | This study | 1C, 1E, 1H, 5B, 6B |
| yKD802 | (1) | lys2::lox-ura3-ISceI ura3-1Δ::KanMx dnl4Δ::HIS3Mx sir3::Nat-GPDp-SIR3 | Batté et al. 2107 | 1C, 6B |
| yKD1562 | (1) | lys2::lox-ura3-ISceI ura3-1Δ::KanMx dnl4Δ::HIS3Mx sir3::Nat-GPDp-SIR3 sae2Δ::HIS3Mx | This study | 1C, 1E, 1H, 6B |
| yKD1474 | (1) | lys2::lox-ura3-ISceI ura3-1Δ::KanMx sae2Δ::HIS3Mx | This study | 1C, 1E, 1H, 5A, 5B, 6A, 6B |
| yKD1656 | (1) | lys2::lox-ura3-ISceI ura3-1Δ::KanMx mre11-H125N | This study | 1C |
| yKD2274 | (1) | lys2::lox-ura3-ISceI ura3-1Δ::KanMx mre11-H125N sae2Δ::Nat | This study | 1C |
| yKD1880 | | MATa RAD5 ADE2 hmlΔ::HPH trp1:: Gal1p-ISceI-TRP1 lys2::lox-ura3-ISceI ura3-1Δ::KanMx Mre11-YFP | This study | 1F, 1G, 7B |
| yKD1881 | | MATa RAD5 ADE2 hmlΔ::HPH trp1:: Gal1p-ISceI-TRP1 lys2::lox-ura3-ISceI ura3-1Δ::KanMx Mre11-YFP sir3::Nat-GPDp-SIR3 | This study | 1F, 1G, 7B |
| yKD1841 | | MATa RAD5 ADE2 hmlΔ::HPH trp1::GAL1p-ISceI-TRP1 lys2::lox-ura3-ISceI ura3-1Δ::KanMx Mre11-YFP sae2Δ::HIS3Mx | This study | 1G |
| yKD1843 | | MATa RAD5 ADE2 hmlΔ::HPH trp1::GAL1p-ISceI-TRP1 lys2::lox-ura3-ISceI ura3-1Δ::KanMx Mre11-YFP sae2Δ::HIS3Mx sir3::Nat-GPDp-Sir3 | This study | 1G |
| yKD1620 | (1) | lys2::lox-ura3-ISceI ura3-1Δ::KanMx sir3::Nat-ADH1p-SIR3 | This study | 1H |
| yKD1845 | | MATalpha RAD5 ADE2 hmlΔ::oripRS hmrΔ::ampR trp1::GAL1p-ISceI-TRP1 TELVI-R::lox-ura3-ISceI ura3-1Δ::KanMx SAE2-GFP-spHIS5 sir3::SIR3-mCherry-HPH | This study | 2A |
| yKD1864 | | MATalpha RAD5 ADE2 hmlΔ::oripRS hmrΔ::ampR trp1::GAL1p-ISceI-TRP1 TELVI-R::lox-ura3-ISceI ura3-1Δ::KanMx SAE2-GFP-spHIS5 sir3::Nat-GPDp-SIR3-mCherry-HPH | This study | 2A, 5C |
| yKD1664 | | MATalpha RAD5 ADE2 hmlΔ::oripRS hmrΔ::ampR trp1:: GAL1p-SceI-TRP1 TELVI-R::lox-ura3-ISceI ura3-1Δ::KanMX | This study | 2B, 2C, 2D, 5E, 6F |
| yKD1680 | | MATalpha RAD5 ADE2 hmlΔ::oripRS hmrΔ::ampR trp1::GAL1p-ISceI-TRP1 TELVI-R::lox-ura3-ISceI ura3-1Δ::KanMx SAE2-GFP-spHIS5 | This study | 2B, 2D, 5E, 7A |
| yKD1712 | | MATalpha RAD5 ADE2 hmlΔ::oripRS hmrΔ::ampR trp1::GAL1p-ISceI-TRP1 TELVI-R::lox-ura3-ISceI ura3-1Δ::KanMx SAE2-GFP-spHIS5 sir3::Nat-GPDp-SIR3 | This study | 2B, 2C, 5E, 7A |

| | | | | |
|---|---|---|---|---|
| yKD1778 | | MATalpha RAD5 ADE2 hmlΔ::oripRS hmrΔ::ampR trp1::GAL1p-ISceI-TRP1 TELVI-R::lox-ura3-ISceI ura3-1Δ::KanMx SAE2-GFP-spHIS5 sir3Δ::Nat | This study | 2B, 2D, 5E |
| Y190 | | MATa his3-Δ200 ade2-101 trp1-901 leu2-3,112 cyh2 gal4Δ gal80Δ ura3-52::URA3::GAL1$_{UAS}$-GAL1$_{TATA}$-lacZ lys2-801::LYS2::GAL4$_{UAS}$-HIS3$_{TATA}$-HIS3 | Harper et al 1993 | 3B,3C, 3D, 4C, 6E |
| yKD1882 | | Y190 sir4Δ::Nat | This study | 3D |
| yKD1991 | | his3-Δ200 ade2-101 trp1-901 leu2-3,112 cyh2 gal4Δ gal80Δ URA3::lexAop-lacZ LYS2::GAL1$_{UAS}$-HIS3$_{TATA}$-HIS3 | This study | 4B |
| yKD2157 | (1) | lys2::lox-ura3-ISceI ura3-1Δ::KanMx sir3::Nat-GPDp-sir3$^{SaiD}$ | This study | 5A |
| yKD2158 | (1) | lys2::lox-ura3-ISceI ura3-1Δ::KanMx sir3::Nat-GPDp-sir3$^{SaiD}$ sae2Δ::HIS3Mx | This study | 5A |
| yKD2161 | (1) | lys2::lox-ura3-ISceI ura3-1Δ::KanMx sir3::Nat-GPDp-sir3$^{SalD-T557I}$ | This study | 5A |
| yKD2162 | (1) | lys2::lox-ura3-ISceI ura3-1Δ::KanMx sir3::Nat-GPDp-sir3$^{SalD-T557I}$ sae2Δ::HIS3Mx | This study | 5A |
| yKD2192 | (1) | lys2::lox-ura3-ISceI ura3-1Δ::KanMx sir3::Nat-GPDp-sir3-T557I | This study | 5B |
| yKD2194 | (1) | lys2::lox-ura3-ISceI ura3-1Δ::KanMx sir3::Nat-GPDp-sir3-T557I sae2Δ::HIS3Mx | This study | 5B |
| yKD2287 | | MATalpha RAD5 ADE2 hmlΔ::oripRS hmrΔ::ampR trp1::GAL1p-ISceI-TRP1 TELVI-R::lox-ura3-ISceI ura3-1Δ::KanMx SAE2-GFP-spHIS5 sir3::Nat-GPDp-sir3-T557I-mCherry-HPH | This study | 5C |
| yKD1932 | (2) | adh4::URA3-TEL7L | This study | 5D |
| yKD1934 | (2) | adh4::URA3-TEL7L sir4Δ::Nat | This study | 5D |
| yKD2181 | (2) | adh4::URA3-TEL7L sir3::Nat-GPDp-SIR3 | This study | 5D |
| yKD2254 | (2) | adh4::URA3-TEL7L sir3::Nat-GPDp-sir3-T557I | This study | 5D |
| yKD2242 | (2) | adh4::URA3-TEL7L sir3-T557I | This study | 5D |
| yKD2286 | | MATalpha RAD5 ADE2 hmlΔ::oripRS hmrΔ::ampR trp1::GAL1p-ISceI-TRP1 TELVI-R::lox-ura3-ISceI ura3-1Δ::KanMx SAE2-GFP-spHIS5 sir3::Nat-GPDp-sir3-T557I | This study | 5E |
| yKD1467 | (1) | lys2::lox-ura3-ISceI ura3-1Δ::KanMx sir3Δ::HIS3MX | This study | 6A, 6B |
| yKD1892 | (1) | lys2::lox-ura3-ISceI ura3-1Δ::KanMx sir4Δ::HIS3Mx | This study | 6A |
| yKD1936 | (1) | lys2::lox-ura3-ISceI ura3-1Δ::KanMx sir3Δ::Nat sir4Δ::HIS3Mx | This study | 6A |
| yKD1969 | (1) | lys2::lox-ura3-ISceI ura3-1Δ::KanMx sir4Δ::HIS3Mx sae2Δ::Nat | This study | 6A |
| yKD2265 | (1) | lys2::lox-ura3-ISceI ura3-1Δ::KanMx sir3-T557I | This study | 6A |
| yKD2278 | (1) | lys2::lox-ura3-ISceI ura3-1Δ::KanMx sir3-T557I sir4Δ::Nat | This study | 6A |
| yKD2038 | (1) | lys2::lox-ura3-ISceI ura3-1Δ::KanMx his3::HIS3-TEFp | This study | 6C |
| yKD2039 | (1) | lys2::lox-ura3-ISceI ura3-1Δ::KanMx his3::HIS3-TEFp-SIR4 | This study | 6C |
| yKD2054 | (1) | lys2::lox-ura3-ISceI ura3-1Δ::KanMx sir3::Nat-ADH1p-SIR3 his3::HIS3-TEFp | This study | 6C |
| yKD2055 | (1) | lys2::lox-ura3-ISceI ura3-1Δ::KanMx sir3::Nat-ADH1p-SIR3 his3::HIS3-TEFp-SIR4 | This study | 6C |
| yKD2042 | (1) | lys2::lox-ura3-ISceI ura3-1Δ::KanMx sir3::Nat-ADH1p-SIR3 sae2Δ::Nat his3::HIS3-TEFp | This study | 6C |
| yKD2043 | (1) | lys2::lox-ura3-ISceI ura3-1Δ::KanMx sir3::Nat-ADH1p-SIR3 sae2Δ::Nat his3::HIS3-TEFp-SIR4 | This study | 6C |
| yKD2133 | (1) | lys2::lox-ura3-ISceI ura3-1Δ::KanMx SAE2-GFP-spHIS5 | This study | 7A, 7C, 7D |
| yKD2138 | (1) | lys2::lox-ura3-ISceI ura3-1Δ::KanMx SAE2-GFP-spHIS5 sir3Δ::Nat | This study | 7C, 7D |
| yKD2139 | (1) | lys2::lox-ura3-ISceI ura3-1Δ::KanMx SAE2-GFP-spHIS5 sir3::Nat-GPDp-SIR3 | This study | 7A, 7C, 7D |
| yKD2141 | (1) | lys2::lox-ura3-ISceI ura3-1Δ::KanMx SAE2-GFP-spHIS5 sir3::Nat-GPDp-SIR3$^{SalD}$ | This study | 7A, 7C, 7D |

| | | | | |
|---|---|---|---|---|
| yKD2334 | | MATa RAD5 ADE2 hmlΔ::oripRS hmrΔ::ampR trp1::GAL1p-ISceI-TRP1 TELVI-R::lox-ura3-ISceI ura3-1Δ::KanMx SAE2-GFP-spHIS5 sir3::Nat-GPDp-SIR3 his3::HIS3-TEFp | This study | 6D, 6F |
| yKD2335 | | MATa RAD5 ADE2 hmlΔ::oripRS hmrΔ::ampR trp1::GAL1p-ISceI-TRP1 TELVI-R::lox-ura3-ISceI ura3-1Δ::KanMx SAE2-GFP-spHIS5 sir3::Nat-GPDp-SIR3 his3::HIS3-TEFp-SIR4 | This study | 6D, 6F |
| yKD2355 | (1) | lys2::lox-ura3-ISceI ura3-1Δ::KanMx sae2-S267E-4ala-YFP | This study | 7E, 7F |
| yKD2356 | (1) | lys2::lox-ura3-ISceI ura3-1Δ::KanMx sae2-S267E-4ala-YFP sir3::Nat-GPDp-SIR3 | This study | 7E, 7F |

(1) All those strains are W303 *MATa RAD5+ ade2-1::ADE2 rap1::GFP-RAP1-LEU2 hml::HPH trp1::GAL1p-ISceI-TRP1*
(2) All those strains are W303 Mata RAD5 ADE2 leu2-3,112 his3-11,15 trp1-1 ura3-1

Bordelet et al, Appendix Table S2

Appendix Table S2**: Primers used in this paper:**

| **Gene** | **Primer Name** | **Primer Sequence** |
|---|---|---|
| OGG1_F | pr776 | CAATGGTGTAGGCCCCAAAG |
| OGG1_R | pr777 | ACGATGCCATCCATGTGAAGT |
| TEL6R_+0.2kb_F | pr750 | TGAGGCCATTTCCGTGTGTA |
| TEL6R_+0.2kb_R | pr751 | CCCAGTCCTCATTTCCATCAA |
| TEL6R_+0.9kb_F | pr752 | TGATGAATTACAAGGGAACAATGAG |
| TEL6R_+0.9kb_R | pr753 | CATCAAACAAGTAGGAATGCGAAA |
| LYS2_0.9kb_F | pr764 | TGATTTACCATTGGGCACAATTT |
| LYS2_0.9kb_R | pr765 | AATTTCCGCGGCAAAGG |
| ISceIcs_F | pr768 | GGAGTTAGTTGAAGCATTAGGTCCC |
| ISceIcs_R | pr769 | GCGGCTTAACTGTGCCCTC |
| LYS2_0.2kb_F | pr760 | GCTCAAAGAGACATGGGTGGA |
| LYS2_0.2kb_R | pr761 | TGTCATCTAAACCCACACCGG |
